# Evolution of SARS-CoV-2 spike trimers towards optimized heparan sulfate cross-linking and inter-chain mobility

**DOI:** 10.1101/2024.07.17.603909

**Authors:** Jurij Froese, Marco Mandalari, Monica Civera, Stefano Elli, Isabel Pagani, Elisa Vicenzi, Itzel Garcia-Monge, Daniele Di Iorio, Saskia Frank, Antonella Bisio, Dominik Lenhart, Rudolf Gruber, Edwin A. Yates, Ralf P. Richter, Marco Guerrini, Seraphine V. Wegner, Kay Grobe

## Abstract

The heparan sulfate (HS)-rich extracellular matrix (ECM) serves as an initial interaction site for the homotrimeric spike (S)-protein of SARS-CoV-2 to facilitate subsequent docking to angiotensin-converting enzyme 2 (ACE2) receptors and cellular infection. Recent variants of concern (VOCs), notably Omicron, have evolved by swapping several amino acids to positively charged residues to enhance the S-protein trimer’s interaction with the negatively charged HS polysaccharide chains in the matrix. These increased interactions, however, may reduce Omicron’s ability to move through the HS-rich ECM to effectively find ACE2 receptors and infect cells, and raise the question of how HS-associated virus movement can be mechanistically explained. In this work, we show that Omicron S-proteins have evolved to balance HS interaction stability and dynamics, resulting in enhanced mobility on an HS-functionalized artificial matrix. Both properties are achieved by the ability of Omicrons S-proteins to cross-link at least two HS chains, providing both high avidity to retain the protein inside the HS-rich matrix, and fast dynamics, thus enabling direct S-protein switching between HS chains as a prerequisite for mobility at the cell surface. Optimized HS interactions can be targeted pharmaceutically, because an HS mimetic significantly suppressed surface binding and cellular infection specifically of the Omicron VOC. These findings suggest a robust way to interfere with SARS-CoV-2 Omicron infection and, potentially, future variants.

## Introduction

Coronavirus disease (COVID-19) is caused by severe acute respiratory syndrome coronavirus 2 (SARS-CoV-2), which first emerged in the city of Wuhan, Hubei province, China, in 2019 [1]. Since then, SARS-CoV-2 has demonstrated unprecedented global spread and rapid evolution into new variants of concern (VOCs). SARS-CoV-2 VOCs bind more rapidly to respiratory epithelial cells, making them more infectious [2]. They also have the ability to evade the immune response generated by previous infections or vaccinations [3]. The Omicron VOC is currently the dominant variant circulating worldwide, and as of January 2024, 99% of the variants circulating in the U.S. were mutations of Omicron, most commonly EG.5 (causing an estimated 24% of all COVID-19), FL 1.5.1 (accounting for 14% of all cases) (https://covid.cdc.gov/covid-data-tracker/#variant-proportions) and, more recently, the KP.2 variant [4]. An important feature acquired during the evolution of the Omicron variants is that they tend to infect the upper respiratory tract (e.g., nose and throat) more than previous variants such as Wuhan and Delta, which tended to infect the lower respiratory tract (e.g., lungs), resulting in more severe disease and higher mortality [5].

SARS-CoV-2 possesses 24 surface spike (S) glycoprotein trimers per virion that are central to host cell binding and infection. Like the S-proteins of SARS coronavirus (SARS-CoV) [6], SARS-CoV-2 S-proteins bind to the cell surface receptor angiotensin converting enzyme 2 (ACE2) [7] (Figure 1a). This S-protein/ACE2 interaction triggers a cascade of events leading to the fusion of cell and viral membranes, facilitating viral entry into host cells. The evolution of this S-protein function is a major determinant of viral infectivity, spread, pathogenesis, and adaptation for infection of new hosts and cell lines [8]. The SARS-CoV-2 S-protein is structurally divided into S1 and S2 subunits. The S1 subunit contains the receptor binding domain (RBD), which is required for binding to ACE2 on the host cell, and the S2 subunit drives subsequent membrane fusion [9]. Importantly, the RBD is hidden within the S1, preventing its interaction with ACE2 [9, 10] and avoiding detection by the immune system. To engage the ACE2 receptor, the RBD must first convert from the inaccessible conformation into an exposed conformation [9, 11] (Figure 1a). This essential conformational change of the S-protein is a consequence of its binding to highly negatively charged heparan sulfate (HS, Figure 1b) polysaccharide chains [11], which are abundant components of the extracellular matrix (ECM) [12].

**Figure 1:**
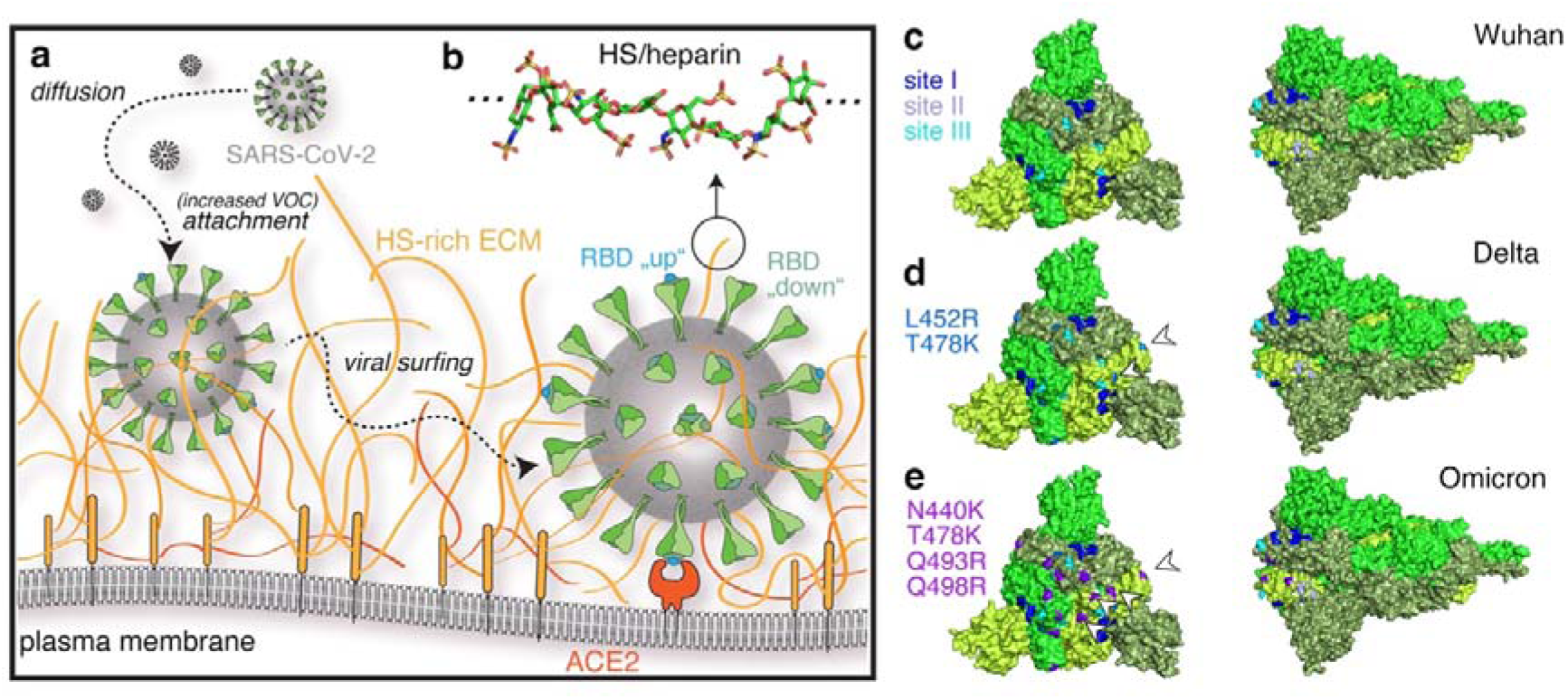
Schematic of SARS-CoV-2 host cell association and invasion. **a)** There are four steps in virus-host interaction: Aerosol inhalation, attachment of viral particles to the ECM, diffusion of viral particles through the ECM (viral surfing), and viral entry through the cell membrane. SARS-CoV-2 attachment to the host cell ACE2 and viral entry require binding of HS (shown in orange) by trimeric S-proteins (green), which bind the virus to the HS-rich ECM environment and causes the RBD subunit of the S-protein to undergo a conformational change to an accessible conformation (from “down”, meaning hidden in the trimeric S1-protein, to the ACE2-accessible “up” conformation shown in cyan) [22]. **b)** HS consists of up to 150 sugar residues, corresponding to chain contour lengths between 25nm and 75 nm (and sometimes even up to 200nm [17, 23]), and is highly negatively charged. Shown is an octasaccharide from heparin (protein data bank (pdb) structure 3irl). Sulfur is shown in yellow, nitrogen in blue, oxygen in red and carbon in green. **c-e)** Mutations on the VOC S-protein (pdb structure 6xr8) to introduce positive charges increase S-protein interactions with negatively charged HS/heparin to improve binding, but are also expected to slow down VOC diffusion (viral surfing) and their ability to find ACE2. Site I amino acid acids R346, N354, R355, K356, R357, and R466 are shown in blue, site II amino acids K424, R454, R457, K458, K462, and R466 are shown in slate, and site III amino acids R403, R408, K417, and K444 are shown in cyan. Acquired amino acids in the Delta VOC are shown in marine blue and indicated by arrowheads, and additional basic residues found in the Omicron BA.1 variant are shown in violet.

In addition to converting the RBD subunit to an accessible conformation for ACE2 binding [11], HS plays a second important role in SARS-CoV-2 infection by acting as a collector of the viruses at the epithelial surface (Figure 1a). Indeed, ACE2 expression levels in human epithelial cells and lung tissue are relatively low, which limits the ability of freely diffusing SARS-CoV-2 to infect cells [13, 14]. To overcome this issue, the virus binds to the HS-rich ECM and moves within it close to the epithelial cell membrane, enhancing the likelihood of interaction with ACE2 [15] (Figure 1a). Given their essential role in cellular infection, the enhanced fitness observed in many SARS-CoV-2 VOCs have therefore been attributed primarily to mutations of the S-protein that dramatically increase both ACE2 and HS binding properties [16] (Figure 1c-e) by increasing the total formal charge of the trimeric S-protein (residues 13-1140) from + 3 (Wuhan) to + 18 and to + 24 for Delta and the Omicron B.1.1.529 subvariant, respectively [17]. These observations suggest that more infectious variants have undergone selection for increased engagement with the negatively charged cell-surface HS. This evolution, however, raises an important question: Do these increased HS interactions impair the rapid movement of SARS-CoV-2 through the ECM and across cell surfaces, a process known as “viral surfing” [18] (Figure 1a) or, do the acquired mutations enhance viral surfing in a counterintuitive manner? If the former were true, the virus would be trapped locally and unable to move to ACE2 receptors on the cell surface. If the latter were true, viral movement on and between cells in tissues would increase and enhance infectivity, as observed for the VOCs. Additionally, important questions include how to mechanistically explain viral surfing of HS-bound viruses and whether the additional positively charged amino acids in Delta and Omicron VOCs enhance the efficacy of strongly polyanionic soluble inhibitors to block S-protein function and cellular infection.

We investigated these questions by using the well-characterized evolution of SARS-CoV-2 VOCs as a model for HS-binding viruses in general. To this end, we compared the interactions of isolated RBDs with sulfated oligosaccharides *in silico*, and the molecular properties of RBDs, trimeric S-proteins and SARS-CoV-2 viral isolates from Wuhan, Delta, and Omicron *in vitro*. We hypothesized that the increased HS binding and decreased HS unbinding of the VOCs [17, 19, 20] would come at the expense of decreased mobility, potentially providing an explanation for the current gradual within-lineage evolution of SARS-CoV-2, and recent basic amino acid reversions (such as Q493R (Figure 1e) to the original glutamine in Omicron BA4 and BA.5), which followed prior rapid emergence of highly divergent lineages. This scenario would have suggested that viral surfing acts as a dynamic selective force against the continued accumulation of positive charge on the S-protein. Alternatively, SARS-CoV-2 Omicron may have reached an optimal state that allows both processes, i.e. strong HS binding and effective movement through the HS-rich ECM, to occur. This latter scenario, however, would require an explanation of the apparent paradox that increased HS interaction strength (i.e, increased *k*_on_ and decreased *k*_off_) does not restrict viral movement to the receptor, as would be expected from short stretches of free virus movement by diffusion interrupted by repeated cycles of prolonged ECM binding and virus immobilization [21].

In this study, we used automated docking analyses to confirm increased free binding energies between the RBDs of the Wuhan, Delta and Omicron variants and a synthetic HS hexasaccharide model, and employed quartz crystal microbalance with dissipation monitoring (QCM-D) in combination with heparin- or HS-functionalized artificial membranes to mimic RBD and S-protein binding to the cellular surface ECM. These methods confirmed increasing HS interactions of the VOC RBDs and trimeric S-proteins, suggesting potentially restricted SARS-CoV-2 mobility on cellular surfaces. However, we also found that trimeric Omicron S-proteins optimized HS binding along with the dynamics of “surfing” on and between HS without the need to detach from HS, potentially allowing this VOC to more effectively scan the cell surface and find the ACE2 receptor. Finally, we observed that the soluble, highly sulfated polysaccharide pentosan polysulfate (PPS) affected S-protein switching *in vitro* and was most effective in preventing Omicron infection in a cell-based bioassay. This suggests that the increased positive charge in Omicron S-proteins and their increased propensity to bind HS and “surf” the chains could be exploited pharmaceutically to target these important first steps in the viral infection pathway.

## Results

### Molecular modelling of VOC S-protein RBD binding to IAGAIA and PPS hexasaccharides reveals the evolution of stronger interactions

HS and heparin polysaccharides consist of alternating N-acetyl D-glucosamine and D-glucuronic acid residues that undergo extensive sulfation and epimerization during biosynthesis (Supplementary Figure 1) [24]. Owing to their high negative charge, linear HS can dynamically bind to positively charged sites on the surface of various soluble proteins [25–27] and viruses [11, 28]. Previous analyses of the Wuhan strain RBD surface revealed three putative heparin-binding sites [22, 29]. Site I is defined by amino acid acids residues R346, N354, R355, K356, R357 and R466 [11], while sites II and III contain K424, R454, R457, K458, K462, R466, and R403, R408, K417, K444, respectively (Figure 1c). A hexasaccharide composed of uronic acid (α-IdoA2S/ β-GlcA) 1-4 linked to N,6-O disulfated glucosamine (α-GlcNS,6S) repeating units with the composition MeO-(4)-α-L-IdoA2S(1→4) α-D-GlcNS,6S (1→4) β-D-GlcA(1→4) α-D-GlcNS,6S (1→4) α-L-IdoA2S α-D-GlcNS,6S(1)-OMe (IAGAIA, Figure 2, Supplementary Figure 1) was previously identified as the minimal interacting heparin epitope [30]. Analogously, a hexasaccharide mimicking pentosan polysulfate (PPS), consisting of repeating, 1→4 linked 2,3-di-sulfo-Xyl-β (14) units, was included in our analysis (Figure 2, Supplementary Figure 1). PPS is related to heparin and HS glycosaminoglycans in terms of anti-inflammatory and anticoagulant properties [31], but has a higher overall negative charge (>3 sulfo groups per disaccharide (PPS) versus <2.4 sulfo groups per disaccharide for heparin and approximately one sulfate group per HS disaccharide [32]). The PPS used here is a plant-derived drug obtained by sulfonation of β-(1→4) xylan from beech wood and is the active ingredient in Elmiron®, an FDA-approved drug for the treatment of interstitial cystitis and for thromboembolic prophylaxis. Furthermore, PPS has recently attracted attention due to interesting activity against SARS-CoV-2 replication [33]. We analyzed the interaction between these two hexasaccharide models and the RBDs of the Wuhan, Delta, and Omicron variants using molecular docking (Glide tool, Schrödinger Inc. [34, 35]) and ranked the selected poses by re-scoring the estimated free energy of binding according to the Molecular Mechanics, General Born Surface Area (MMGBSA) [36] approach, a method to calculate free energies of ligand-protein interactions (Prime tool, Schrödinger Inc. [37]).

**Figure 2:**
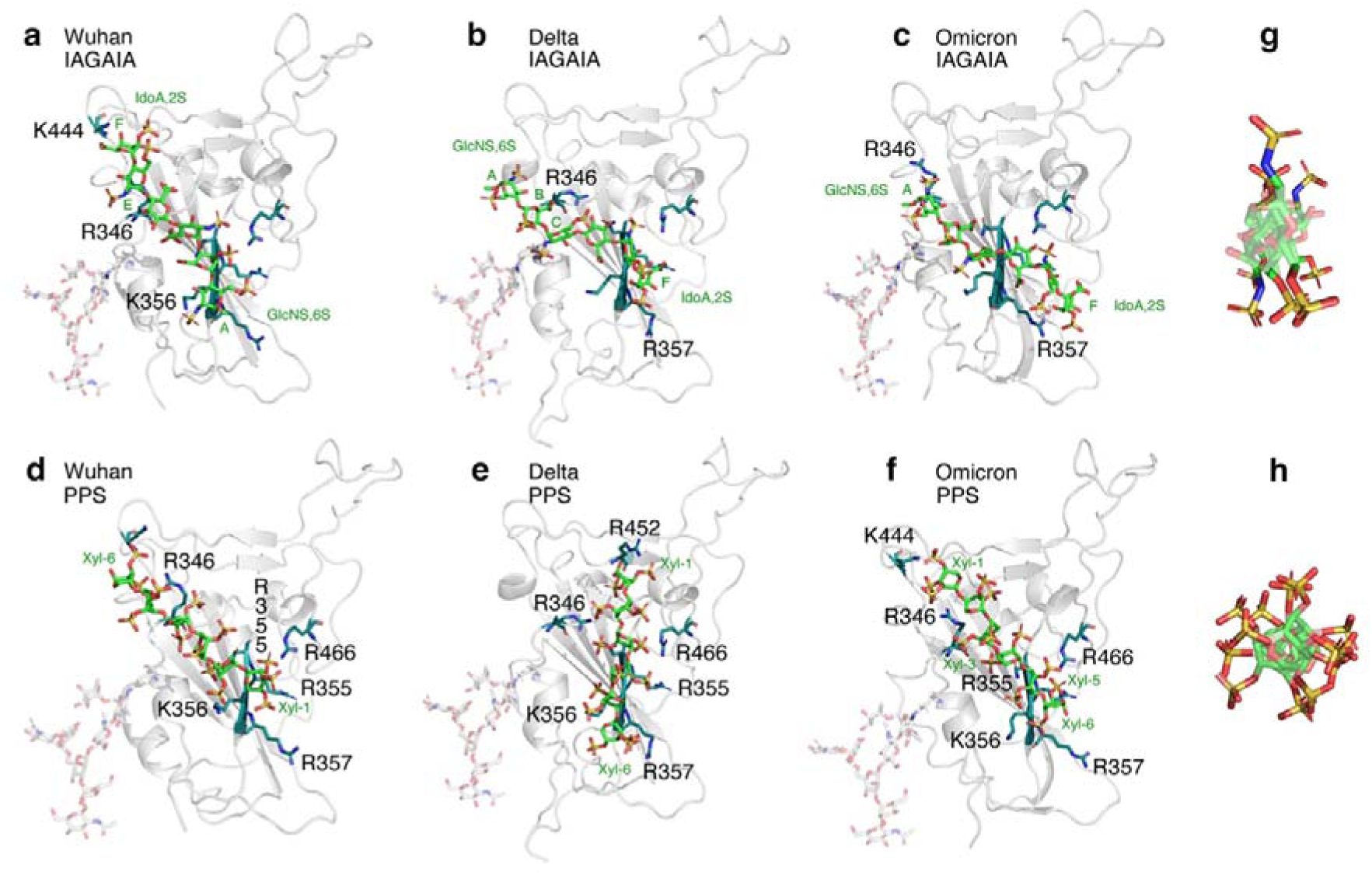
**a-f)** Top-ranked pose of IAGAIA (a-c) [30] and PPS (d-f) in the RBD of the SARS-CoV-2 Wuhan (PDB ID: 6M0J), Delta (PDB ID: 7V8B) and Omicron (PDB ID: 7WBP) VOCs. The S-protein RBD region is represented by a grey cartoon, while IAGAIA and PPS are represented by green, red, blue and yellow sticks indicating carbon, oxygen, nitrogen and sulfur atoms, respectively. The key conserved residues of the pocket (R346, N354 R356, K356, R357, K444 and R466) and the L452R mutation of the Delta RBD in PPS docking are depicted with similar colour codes (deep teal for carbon, red for oxygen and blue for nitrogen). The interaction between these two hexasaccharide models and the RBDs of Wuhan, Delta, and Omicron variants has been characterized using molecular docking (Glide tool, Schrödinger Inc. [34, 35]), while the ranking of the selected poses has been re-scored estimating the free energy of binding according to the MMGBSA [36] approach (Prime tool, Schrödinger Inc. [37]). See Supplementary Tables 1-2 for details. **g,h)** The transversal section of IAGAIA and PPS shows an interesting structural feature that differentiates both ligands: while IAGAIA orients the sulfo-groups linearly to form two opposite negatively charged sides, PPS has a more radial distribution of sulfates, in turn generating a uniform negatively charged surface.

Molecular docking predicts that glycosaminoglycan oligosaccharides IAGAIA and PPS preferentially bind site I of the Wuhan RBD: residues R346, N354, R355, K356, R357 and R466 are found within 3 Å from the oligosaccharide in the best-selected poses (Supplementary Table 3), despite their different relative positioning and orientation (Figure 2, Supplementary Figure 2a-c). Indeed, the best pose between IAGAIA and the Wuhan RBD shows close contact between residues of site I and monosaccharide sugars GlcNS6S (E in Supplementary Figure 1), GlcNS6S (C), IdoA (B), and GlcNS6S (A), in that order (Figure 2a). In contrast, the best pose of IAGAIA and the Delta RBD shows that this set of amino acids contacts the following sugars: IdoA (B), GlcNS6S (C), IdoA (D), GlcNS6S (E), and IdoA (F), in that order (Figure 2b). The *head-tail* orientation of IAGAIA in the Delta RBD is opposite to that predicted in the Wuhan RBD model. Even though the total number of contacts between IAGAIA and site I do not significantly change among both RBD models, the number of salt bridges and van der Waals contacts increases, and correspondingly, the estimated free energy of binding increases (slightly) from -18 to -26 Kcal/mol (Table 1). The IAGAIA in the bound state with the Omicron RBD shows an epitope binding comparable to that of the Delta model and reveals the same monosaccharide – protein contact order: residues R346, N354, R355, K356, R357 and R466 are in contact with the monosaccharides GlcNS6S (A), IdoA (B), GlcNS6S (C), IdoA (D), GlcNS6S (E), and IdoA (F), in that order (Figure 2 c). The estimated free energy of binding that characterizes the interaction between IAGAIA and Omicron is significantly greater than what we observed when the same glycan binds the Delta and Wuhan RBDs, as shown by the higher number of favourable contacts compared to the other models. This suggests a greater propensity of the surface of Omicron RBD to host heparin oligosaccharides (Table 1).

**Table 1.** Average MMGBSA re-scoring valuess, salt bridges, hydrogen bonds and overall contacts of the 10 poses obtained for SARS-CoV-2 VOCs RBDs and IAGAIA/PPS ligands.

|  |  | Free binding energy (kcal/mol) | Salt Bridges | H-bonds | Total contacts |
| --- | --- | --- | --- | --- | --- |
| IAGAIA | Wuhan (site I) | -18 | 1 | 2 | 5 |
|  | Delta (site I) | -26±9 | 5 | 1 | 8 |
|  | Omicron (Top-ranked) | -37 | 5 | 5 | 7 |
|  | Omicron (site I) | -31±5 | 6 | 5 | 16 |
|  | Omicron (site II) | -33±2 | 4 | 3 | 9 |
|  |  | Free binding energy (kcal/mol) | Salt Bridges | H-bonds | Total contacts |
| PPS | Wuhan site I | -24 | 5 | 4 | 11 |
|  | Wuhan site II | -24±3 | 6 | 3 | 11 |
|  | Delta (site I) | -27±5 | 5 | 6 | 15 |
|  | Omicron (site I cluster a) | -31±5 | 6 | 4 | 18 |
|  | Omicron (site I cluster b) | -24 | 4 | 5 | 18 |

The PPS oligosaccharide displays a radial distribution of negatively charged sulfated groups along its molecular axis (Figure2 g,h). Therefore, we propose a higher probability of the PPS sulfate groups to engage, in non-directional salt bridge interactions, the positively charged patches of Arg and Lys residues that are solvent exposed on the RBD surface (Supplementary Figure 2d-f). In the best docking pose of PPS in the Wuhan RBD model, the monosaccharide units Xyl-6, Xyl-2, and Xyl-1, in that order, are closer to site I residues (Figure 2 d). Interestingly, PPS shows a greater number of salt bridges and contacts, probably correlated to both an increase of the negative charges and a uniform distribution of sulfate groups compared to IAGAIA. This allows for more efficient interactions with the basic channel of the RBD surface (Supplementary Table 3). Interestingly, the contact efficiency between PPS and RBD increases from Wuhan to Delta to Omicron RBD variant models, respectively, and the free energy of binding also increases (Table 1). Indeed, in the best pose of PPS in both the Delta and Omicron RBDs, all residues of site I engage the ligand at the Xyl-5, Xyl-4, Xyl-2 and Xyl-1 units via an additional contact between the mutated residue R452 and Xyl-1 (Figure 2e).

In summary, the docking results for the IAGAIA and PPS oligosaccharides show a preference for binding to site I for all of the VOC RBDs. The analysis of the docking calculations suggests that the interactions between the basic RBD residues and the negatively charged groups of IAGAIA and PPS are mainly charge-based. For PPS, the docking results reveal a conserved binding mode for each RBD tested, establishing an extended network of salt bridges and H-bonds compared to IAGAIA. This increases the affinity to sulfated glycans during the evolution of Wuhan to Delta and to Omicron RBDs (Table 1, Supplementary Table 2).

### SARS-CoV-2 VOC S-protein RBDs have evolved towards increased HS crosslinking

Next, we tested the prediction of improved glycan interactions during SARS-CoV-2 evolution with HS [17, 19, 38] using QCM-D. The core of QCM-D is an oscillating quartz crystal sensor disk with a resonance frequency related to the mass of the disk (Supplementary Figure 4a). This allows the real-time detection of nanoscale mass changes on the sensor surface by monitoring changes in the (normalized) resonance frequency (Δ*F*): adsorption of molecules on the surface decreases *F*, while mass loss of mass increases *F* (Supplementary Figure 4b). QCM-D measures an additional parameter, the change in energy dissipation *D*, which is particularly useful for studying soft layer properties, as Δ*D* correlates with layer softness (Supplementary Figure 4b).

Using this technique, we investigated the interaction of RBDs with HS functionalized surfaces, which were formed on top of supported lipid bilayers (SLBs) as a plasma membrane model (Supplementary Figure 4a). We used heparin instead of HS because the RBD, monomeric S-protein, and trimeric S-proteins show strong binding to polysaccharides that are composed of trisulfated repeating units (IdoA2S-GlcNS6S) [30], a motif typically found in heparin chains. The SLBs were formed through the rupture of small unilamellar vesicles (SUVs) containing 5 mol% biotinylated lipids. Highly sulfated heparin chains were anchored to the SLBs through a streptavidin linker *via* a biotin moiety at the reducing end and served as a proxy for cell surface HS (Supplementary Figure 4a). To mimic RBD contact with the extracellular matrix, we then added the RBD and monitored mass increases upon its adsorption to the heparin surface, as indicated by decreased resonance frequencies (-Δ*F*), as well as changes in the softness of the heparin film upon protein binding (as reflected by the dissipation shift Δ*D*).

As shown in Figure 3a, addition of unlabeled RBDs of the Wuhan, Delta and Omicron variants on top of a heparin functionalized QCM-D chip induced rapid frequency decreases (-Δ*F*) of approximately 60 Hz, indicating similarly fast binding of all RBDs to heparin (and specific binding to heparin and not the lipid or streptavidin, Supplementary Figure 4c). We also observed that the energy dissipation (Δ*D*) of the layer decreased proportionally during RBD binding, suggesting protein-induced stiffening of the heparin film, which could be caused by cross-linking the heparin chains. However, while the overall decrease in *D* was very similar for all RBDs (Figure 3a, black asterisk), indicating similar capacities to cross-link surface heparin, we found that the decrease in *D* occurred earlier (that is, at a lower RBD surface coverage) for Omicron and Delta RBDs compared to the Wuhan RBD. This finding suggests a higher cross-linking propensity of these variants compared to the Wuhan RBD (red asterisk). Parametric plots of Δ*D*/-Δ*F* (a measure of film softness) vs. -Δ*F* (a measure of surface density) confirmed the evolving cross-linking propensity (Figure 3b), with more pronounced rigidification (and thus cross-linking) of the Delta and Omicron RBDs compared to the Wuhan RBD at any given RBS surface coverage (-Δ*F* shift), except at the highest coverage where the degree of stiffening is similar for all variants. The same effect was also observed when a lower-sulfated HS was coupled to the QCM-D sensor surface instead of heparin (Figure 3c). This confirmed that SARS-CoV-2 RBD binding and the associated but distinct HS film stiffening are qualitatively preserved over different HS sulfation levels.

**Figure 3:**
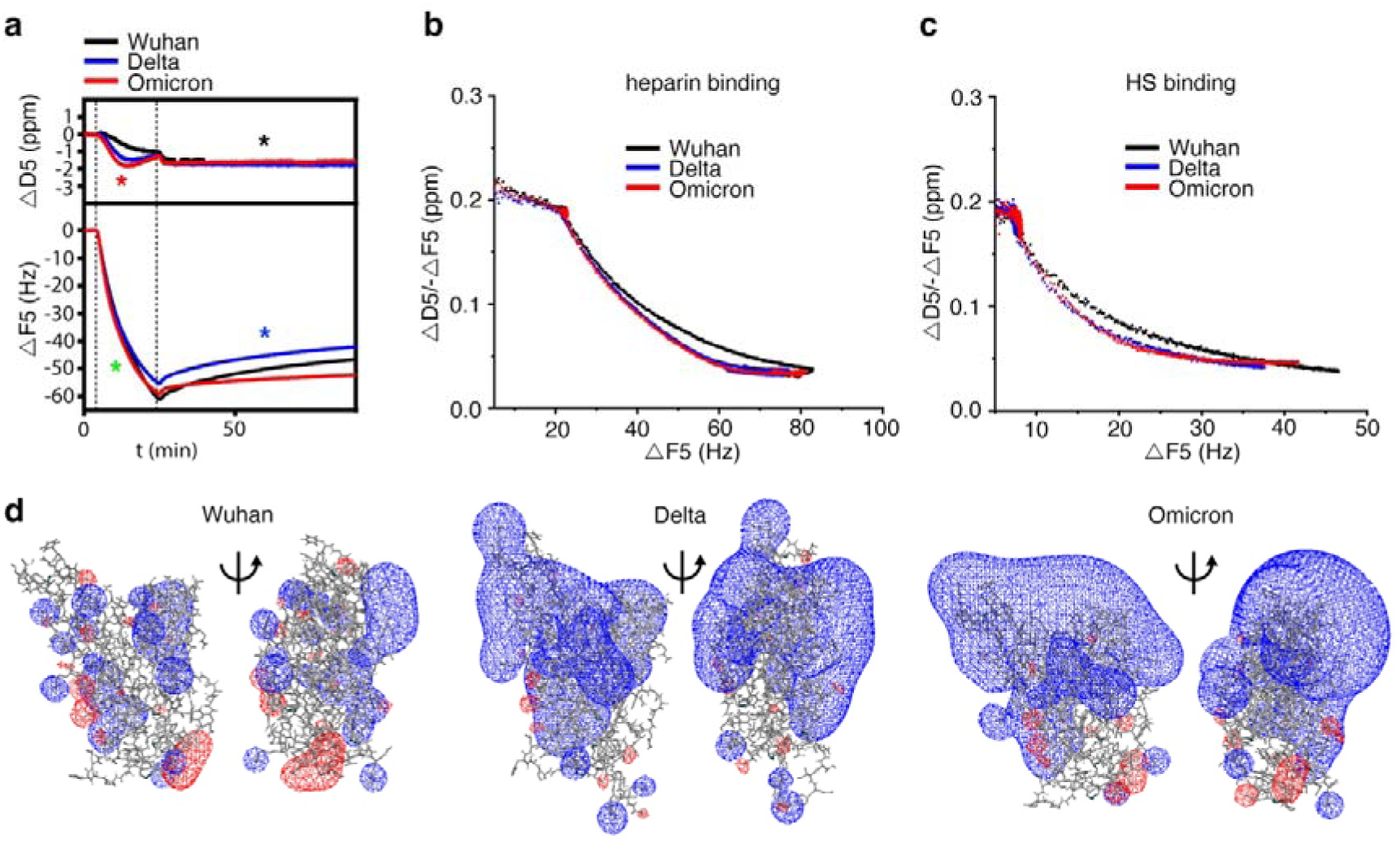
Delta and Omicron RBDs evolved towards increased heparin cross-linking and stability of the heparin interaction. **a)** Unlabeled RBD binding kinetics to heparin are similar (green asterisk), as indicated by similar Δ*F* decrease profiles during RBD binding, and the ability of VOC RBDs to cross-link heparin are also similar (black asterisk). In contrast, Delta and Omicron RBDs improved their potency to cross-link heparin (red asterisk) and the stability of the interaction (blue asterisk). **b,c)** Parametric plots of Δ*D*/-Δ*F* (a measure of film softness) vs. -Δ*F* (a measure of surface density) confirmed very similar heparin (b) / HS (c, Supplementary Figure 4 d-f) cross-linking abilities for Delta and Omicron RBDs and reduced heparin/HS cross-linking ability of the Wuhan RBD. **d)** Comparison of electrostatic potentials for Wuhan (*Left*), Delta (*Center*), and Omicron (*Right*) RBD variants. The electrostatic potential was visualized using the SWISS-MODEL [39]. Color representation: Blue: Positive charge, Red: Negative charge.

Finally, we observed that while all RBDs remained largely bound during wash buffer injection into the QCM-D chambers, dissociation of the Wuhan RBD (and to a lesser degree also the Delta variant) was increased relative to that of the Omicron variant, as evidenced by an observable increase in Δ*F* during the 60-minute wash step (Figure 3a, blue asterisk, Supplementary Figure 4d-f). Taken together, the faster increase in heparin layer stiffness (evidenced by -Δ*D*) and the increased half-life of the interaction (evidenced by small changes in Δ*F* during prolonged washing of the sensor surface) indicate that the Omicron RBD has evolved to rapidly establish multivalency and to reduce the *k*_off_ of the interaction. These findings are supported by the *in silico* calculations predicting stronger HS binding of the Delta and Omicron RBDs (Supplementary Table 1) and by an increase of the number of contacts (Figure 2, Supplementary Table 3). Nevertheless, the RBD association kinetics with the heparin-functionalized sensor surface are not much affected by accumulated charge during evolution, as evidenced by a similar slope of Δ*F* during RBD binding (Figure 3a, green asterisk). Instead, these evolutionary changes ensure that the virus remains bound to HS more effectively by increasing the cross-linking kinetics so that the newer virus variants do not diffuse away from the cell surface, which would hinder infection. Finally, we note that despite the relatively short range of electrostatic forces of soluble proteins at physiological ionic strength, their marked increase in the RBDs of the newer virus variants (Figure 3d) may facilitate favorable RBD orientations toward HS of the opposite charge to increase the speed of HS/heparin cross-linking over short distances.

### SARS-CoV-2 VOC S-protein RBDs have evolved towards increased HS interaction stability

Although our results indicated only modest rates of dissociation of VOC RBDs from the sensor surface into the wash buffer, this is true in general for surface measurements due to analyte rebinding. In these cases, the exchange rate between surface molecules can be obtained by the addition of soluble acceptor ligands. To account for this, we adapted our previously established protocol [40] and injected increasing amounts of soluble heparin onto the RBD-loaded heparin film as a competitor.

As shown in Figure 4a and Supplementary Figure 5a, soluble heparin served as a moderately effective acceptor for the RBD of the Wuhan variant, as evidenced by an increase in *D* (black top asterisk) and *F* as a result of RBD desorption from the sensor surface into the soluble phase (black bottom asterisk, and Supplementary Figure 5b). We also found that soluble heparin desorbed the Delta RBD, at least at high heparin concentrations. In remarkable contrast, the RBD of the Omicron variant remained firmly associated with the QCM-D sensor surface, even when challenged with 12 mM heparin as a soluble acceptor (Figure 4a, red asterisks). This indicated that Omicron RBD has slower disassociation kinetics from HS chains compared to the Wuhan variant RBD, consistent with no observable dissociation during buffer wash (Figure 3a, Supplementary Figure 5b). Furthermore, we observed the same trend when PPS was added to the wash buffer instead of heparin (Figure 4b, and Supplementary Figure 5c) [41]. Consistent with the increase in negative charge, the capacity of PPS to elute the RBDs of the Wuhan and Delta variants was higher than that of heparin. However, the RBD of the Omicron variant also remained firmly associated with the QCM-D sensor surface when challenged with PPS (Figure 4b, red asterisk, Supplementary Figure 5d). This confirmed that the Omicron RBD has evolved towards more persistent HS binding, probably facilitated by its ability to bind two or more chains simultaneously and the increased free binding energies of interaction (Table 1).

**Figure 4:**
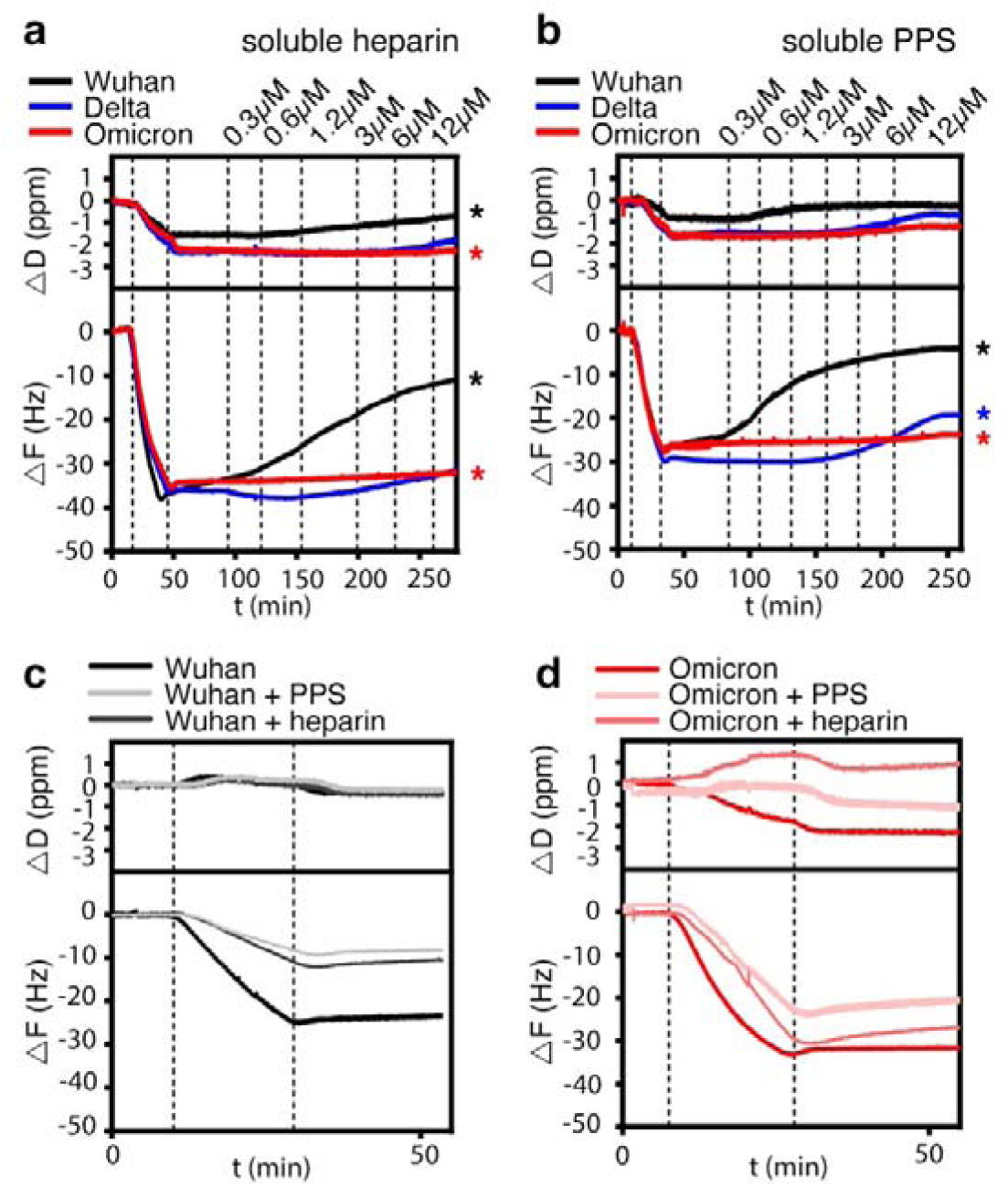
Soluble heparin and PPS increase Wuhan and Delta RBD desorption, but not that of the Omicron RBD. **a)** The dissociation of the Wuhan RBD from the sensor surface increases with increasing heparin concentration in the washing buffer (black asterisk), and the Delta RBD also dissociates at higher heparin concentrations. The Omicron RBD does not dissociate from the heparin-functionalized sensor surface (red asterisk). **b)** Increased negative charge of soluble PPS added to the wash buffer increases dissociation of the Wuhan and Delta RBDs (black and blue asterisks), but again not of the Omicron RBD (red asterisk). **c)** Preincubation of the Wuhan RBD with 100mM heparin or PPS inhibit S-protein association with the sensor surface to a similar extent. **d)** In contrast, preincubation of the Omicron RBD with heparin inhibits its association with the sensor surface to a lesser extent than PPS, which is also only moderately active.

It has been shown previously that pre-incubation with soluble heparin impairs subsequent S-protein or RBD binding to cells and abrogates SARS-CoV-2 infection [11, 15, 29, 30, 42, 43]. As an *in vitro* model, we preincubated the RBDs with 100 nM heparin or PPS before passing them over the heparin-functionalized QCM-D sensors. Pre-incubation of 200nM (5µg/ml) of the Wuhan variant RBD with heparin or PPS greatly delayed protein association with the heparin-functionalized sensor surface, as evidenced by a strongly reduced binding slope and a strongly reduced –Δ*F* (Figure 4c). Therefore, preincubation with heparin or PPS is a potential pharmaceutical approach to reduce cellular infection by decreasing the likelihood of virus binding to the cells [44, 45]. The largely unchanged Δ*D* suggests that the Wuhan RBD does not stiffen the matrix much, presumably due to the reduced cross-linking of the preincubated RBD, as it is already glycan-associated. In contrast, heparin and PPS preincubation with Omicron RBD altered the surface-binding to a lesser extent, and the capacity of the pretreated Omicron RBD to cross-link surface-bound heparin was strongly reduced (Figure 4d). These results suggest that PPS preincubation inhibits the association of the Omicron RBD with the sensor surface to a greater extent than heparin, which is only moderately effective, while both heparin and PPS more effectively inhibit the association of the Wuhan variant RBD. These results are consistent with a previous study showing that PPS inhibits most S-protein RBD mutants better than heparin [46]. The results also support the idea that the S-protein RBD does not recognize a specific HS binding motif, and relies on non-selective electrostatic interactions with the HS in the ECM. This, in turn, highlights the importance of increasing charge differences between the S-proteins and the ECM during viral evolution to maintain the initial contact and to enhance the chances of subsequent cellular infection.

### Trimeric SARS-CoV-2 VOC S-proteins have evolved towards increased capability for intersegmental transfer between HS

The results so far supported the contention that negatively charged glycans, particularly HS, act as attachment factors for SARS-CoV2 to the ECM, and that maintaining the interaction has been optimized during the evolution of the Omicron RBD [17, 19]. However, diffusion of SARS-CoV-2 through the mucous layer of the airway epithelium (viral surfing) is also essential for binding to ACE2, cellular entry, and infection [47]. In this regard, our results suggest a detrimental effect of increased HS interaction half-life, since the newer SARS-CoV-2 variants are more likely to be trapped in the ECM. This would reduce their viral surfing capabilities by intermittent free diffusion, which would stand in contradiction to the observed increase in infectivity and transmissibility of the VOCs. We hypothesized that this discrepancy may possibly be explained by different HS binding of the VOC RBDs and the trimeric S-proteins that decorate the viral surface. In fact, the avidity to HS is likely to be higher for the trimeric S-protein than for the monomeric RBD, due to the cooperative effect provided by its multimeric state (Figure 1c-e)[30, 48, 49], and is supported by the observation that for the Wuhan variant the RBD domain binds to heparin with moderate affinity (*K*_D_ ∼1 μM), whereas the trimeric S-protein binds with much higher affinity (*K*_D_ = 64 nM) [30]. We also observed that the acquired mutations of the trimeric Omicron S-proteins make a sizeable contribution to the affinity of the protein/heparin interaction, decreasing the dissociation constant (*K*_d_) from 2.65 ± 0.4 nM for the Wuhan variant to 1.5 ± 0.2 nM for the Omicron variant, as determined by the established QCM-D set-up (Supplementary Figure 6a). However, we again note that such increased S-protein interactions with negatively charged HS would be expected to interfere with, rather than enhance, the rapid movement of SARS-CoV-2 Omicron through the ECM and across the surfaces of epithelial cells.

To resolve the potential paradox of increased binding affinity increasing viral surfing, we investigated the binding of trimeric S-proteins of the Wuhan and Omicron variants to the heparin functionalized QCM-D chips [40], followed by injection of increasing amounts of soluble heparin. As shown in Figure 5a and Supplementary Figure 6b and 6d, soluble heparin served as a moderately effective acceptor for the Wuhan variant trimeric S-proteins, and a large fraction of the protein could not be displaced from the surface even at high soluble heparin concentrations. In remarkable contrast to the Wuhan trimeric S-protein and the Omicron RBD in the same experimental setup, the trimeric Omicron S-proteins desorbed much faster from the QCM-D sensor surface, even at low heparin concentrations of 0.3 μM (Figure 5a, Supplementary Figure 6c and 6e). Overall, a larger fraction of the Omicron trimeric S-protein could be displaced by heparin than for the Wuhan variant, despite the opposite behavior of the respective monomeric RBDs.

**Figure 5:**
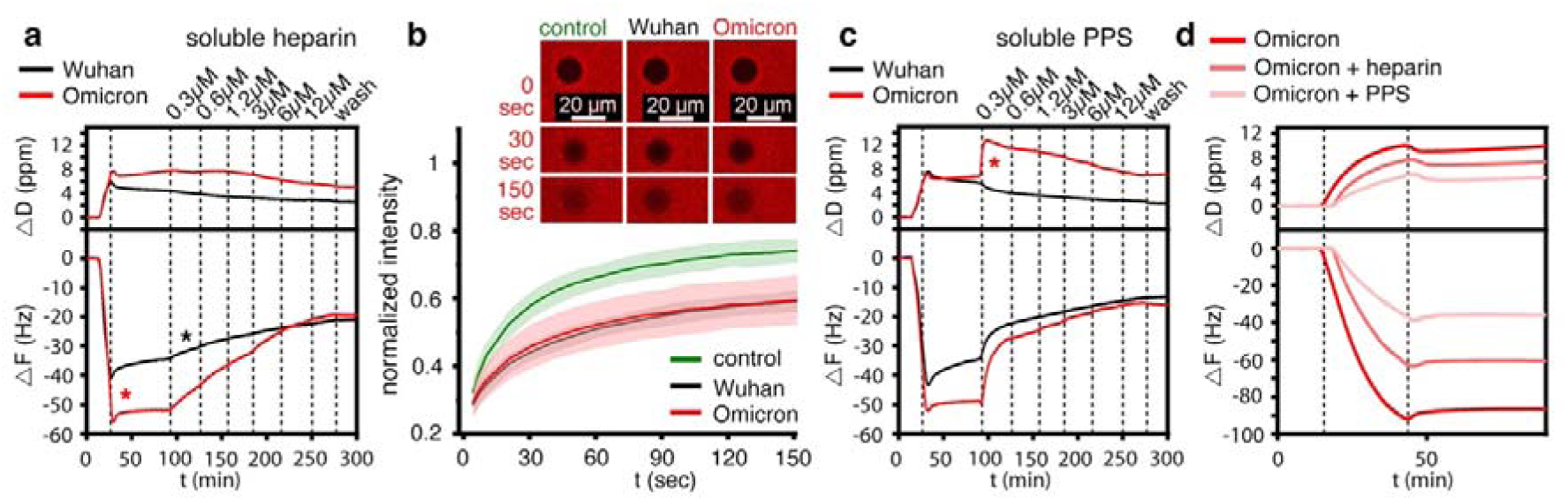
Homo-trimerized S-proteins show drastic changes in binding characteristics to HS. **a)** The relative amount of trimeric Omicron S-protein binding to the heparin film is increased over that of the trimeric Wuhan S-protein (red asterisk), despite similar protein concentrations in the loading buffer and similar loading time. During washing without soluble competitor (25 to 90 min), the dissociation of the trimeric Wuhan S-protein was increased over the trimeric Omicron S-protein. The trimeric Omicron S-protein remained tightly bound to the sensor surface, as previously observed for the Omicron RBD. Binding of both trimeric S-proteins resulted in an increase in dissipation. Addition of increasing amounts of soluble heparin (starting from 90 min) increases the desorption of the trimeric Wuhan S-protein (black asterisk) from the sensor surface much less than the desorption of the trimeric Omicron S-protein, as indicated by the different slopes of the decreasing -Δ*F*. **b)** FRAP confirmed similar cross-linking of heparin chains by trimeric Wuhan and Omicron S-proteins, as evidenced by similarly decreased mobility of fluorescently labeled heparin/streptavidin on the functionalized sensor surface. Top: Representative fluorescence micrographs displaying the FRAP results. Shown are bleach areas at 0 sec and post-bleach areas at 30 sec and 150 sec. control: no added protein. **c)** Increased negative charge of soluble PPS added to the wash buffer increases the desorption of the trimeric Omicron S-proteins the most. Note the increase in *D*, possibly due to PPS-induced de-crosslinking of heparin chains (red asterisk). **d)** Preincubation of trimeric Omicron S-proteins with heparin or PPS strongly inhibits their subsequent binding to surface-bound heparin, and the inhibition of the trimeric Omicron S-proteins with PPS is more effective than with heparin.

A second important observation was that the association of the trimeric Omicron S-protein with the heparin film on the sensor was more effective, as evidenced by the decreased *K_d_* (Supplementary Figure 6a) and by the larger -Δ*F* of the trimeric Omicron S-protein compared to that of the Wuhan variant (20% more protein binding), despite the same concentrations and loading times used for both proteins (Figure 5a). Possibly, this observation can be explained by increased electrostatic potential of the Omicron S-protein that attracts the diffusing protein close to the heparin layer prior to binding (a process called “electrostatic steering”) [17]. Finally, trimeric S-protein association increased Δ*D*, which may have resulted from the larger molecular size of the trimers (410.1 kDa) if compared to the monomeric RBDs (26.5 kDa). Indeed, complementary fluorescence recovery after photobleaching (FRAP) assays suggest that the RBDs and the trimeric S-proteins share the ability to cross-link heparin, as evidenced by a reduced 2D diffusion rate of fluorescently labeled heparin-coupled streptavidin on the SLB by the trimeric S-proteins (Figure 5b).

Next, we passed increasing amounts of soluble PPS on top of the heparin-associated trimeric S-proteins of the Wuhan and Omicron variants. As shown in Figure 5c, soluble PPS served as a more effective acceptor of the trimeric Wuhan and Omicron S-proteins than heparin at the same concentration, leading to much faster detachment from the sensor surface (Supplementary Figure 6b and 6d). Notably, even the lowest concentrations of PPS rapidly increased *F* as a result of trimeric S-protein desorption from the sensor surface and its association with the highly negatively charged soluble acceptor. However, we also found that trimeric S-protein desorption remained incomplete even in the presence of increased amounts of PPS, as observed before for heparin (Figure 5a). This result suggests that the trimeric S-protein associates with the heparin film in two distinct modes, one mode allowing direct switching to soluble acceptors and the other mode representing less reversible binding to the sensor surface. We found that the evolution of the trimeric Omicron S-protein promoted the former binding mode over the latter (more static) binding mode, as the desorption of the trimeric Omicron S-protein by soluble PPS was accelerated more than that of the trimeric Wuhan S-protein (Figure 5c, Supplementary Figure 6c and 6e). Taken together, trimeric VOC S-proteins have evolved to improve their ability to directly switch between HS chains, i.e. to surf the chains without the need to first detach from them, despite their isolated RBDs evolving in the opposite direction - possibly because there is no evolutionary pressure on the RBD alone, only on the functional surface trimer. We propose that this allows for persistent ECM binding of trimeric S-proteins and their ability to move within the ECM, allowing the virus to effectively search the cell surface for the ACE2 receptor.

In support of this concept, we also found that preincubation of the trimeric Omicron S-protein with 100 nM heparin or PPS greatly delayed its association with the heparin functionalized sensor surface, as evidenced by a greatly reduced -Δ*F.* As shown in Figure 5d, preincubation with PPS inhibited the binding of trimeric Omicron S-proteins to surface-bound heparin more than heparin, and the effect was stronger than that seen for the trimeric Wuhan S-proteins (Supplementary Figure 6f and 6g). Inspired by these results, suggesting that soluble PPS may have the potential to desorb SARS-CoV-2 from the ECM (Figure 5c) and then remain associated with the viral surface to inhibit rebinding to cellular surfaces (Figure 5d), we tested next whether PPS treatment is a potential pharmaceutical approach to reduce cellular infection specifically by the Omicron VOC [44, 45].

### PPS is the most potent inhibitor of Omicron in SARS-CoV-2 plaque-forming assays

We have previously shown that both heparin and PPS inhibit SARS-CoV-2 infection in Vero cells [29, 33]. However, these studies were performed with the very first variant of SARS-CoV-2 that was predominant in Europe in early 2020 (this strain is characterized by a D614G mutation) in the S-protein [50]. We have now extended the study of heparin and PPS activity against the Omicron variant by testing both compounds side-by-side in a plaque reduction assay. For this purpose, heparin and PPS were tested against a fixed amount of virus, i.e. 50 plaque forming units (PFU), previously titered in Vero cells. As shown in Figure 6a and 6d, heparin, and, even more so, PPS, significantly inhibited D614G and Omicron plaque formation by reducing the number of plaques to 50% or more when cells were pretreated. The same effect was reproduced by pre-treating the virus with both heparin and PPS (Figure 6b and 6e). However, treating both the virus and cells (cell + virus pre-treatment) with either heparin or PPS was more efficient than single treatment, with PPS reducing plaque formation to almost 90% (Figure 6c and 6f). These results show that the higher number of mutations in the RBD of the Omicron S-protein than D614G [51] maintain or even increase the inhibitory effect of heparin and PPS on viral infection; PPS showing highest efficiency in the inhibition of cellular infection by the Omicron BA.1 variant.

**Figure 6:**
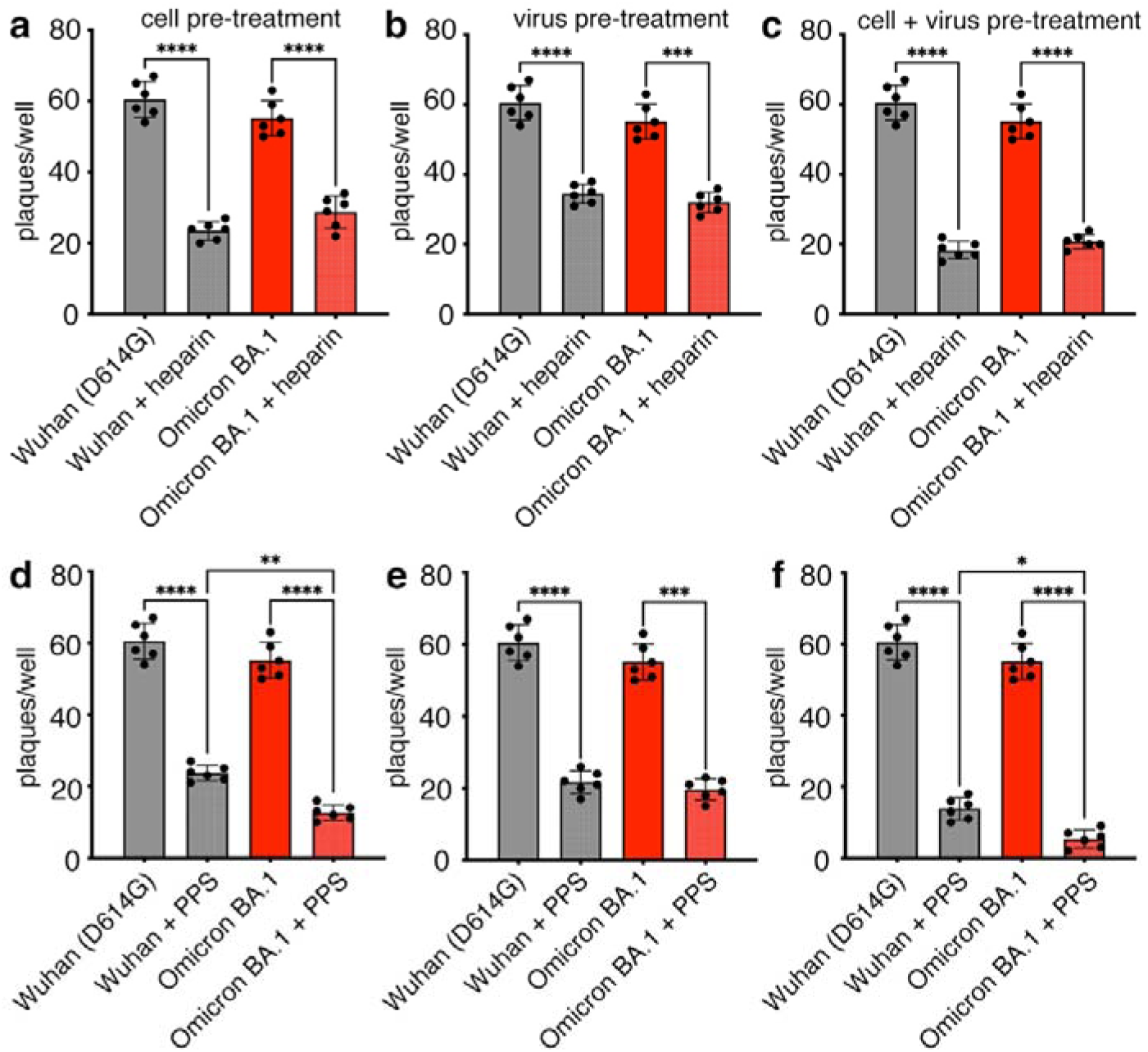
Antiviral activity of heparin and PPS. Plaque formation in Vero cells was determined after infection with either the D614G or Omicron BA.1 variants. Either cells (cell pre-treatment, **a,d**) or viruses (virus pre-treatment, **b,e**) or both (cell + virus pre-treatment, **c,f**) were treated with 100 µg/ml heparin and PPS. Bars represent the mean ± SD of two independent experiments performed in triplicate. Comparisons between groups were made using the one-way analysis of variance (ANOVA). The mean of each treatment column was compared with the mean of each other column. Tukey correction was used for multiple comparisons. ****: p<0.0001; ***: p<0.001; *: p=0.038. See Supplementary Table 5 for details.

## Discussion

Rapid and specific interactions between the SARS-CoV-2 S-protein and largely negatively charged ACE2 receptors are fundamental to cellular infection. It is therefore not surprising that the S-protein, and in particular the RBD, showed a steady increase in positive charge to increase binding that was remarkably accelerated when the Delta variant was leading the pandemic [38, 52]. It was quickly found that with this increase in charge, the binding affinity between spike VOCs and the negatively charged cell surface HS increased as well [17, 19, 38]. Remapping the positive charge and optimizing the electrostatic S-protein interactions with both HS and ACE2 was therefore a major driver in the evolution of SARS-CoV-2 into one of the most infectious viruses in human history [53].

In our experimental work, we used QCM-D to characterize the effect of VOC RBD and trimeric S-protein evolution on the binding of the cellular glycocalyx using a heparin-functionalized lipid bilayer as a proxy [40, 54, 55]. The first important result of our QCM-D analyses is that the RBDs of the Wuhan, Delta and Omicron variants share the ability to cross-link at least two heparin chains on the functionalized surface, and the RBDs of the latest SARS-CoV-2 variants have a stronger propensity to cross-link heparin. Consistent with this, mean binding energies with a hexasaccharide of -18 kcal/mol (Wuhan) increased to -27 kcal/mol (Delta) and to -37 kcal/mol (Omicron). We show that one consequence of these evolved properties is a reduction in the off-rate of the RBD/HS interaction, as it is mediated by multivalent interactions at multiple sites that are unlikely to unbind simultaneously (a process called the multivalent effect, Figure 7a). In the *in vivo* situation, such evolutionary optimization of multivalent HS binding could translate into more stable interactions, confining the virus to the site of initial ECM contact and reduce its spread by fluid flow and diffusion to more distant sites. This, in turn, may be a contributing factor to the altered pathophysiology of SARS-CoV-2, which began as lower respiratory tract infections with lower infectivity in the earlier variants and developed into a predominantly upper respiratory tract infection with higher infectivity in the later variants.

**Figure 7:**
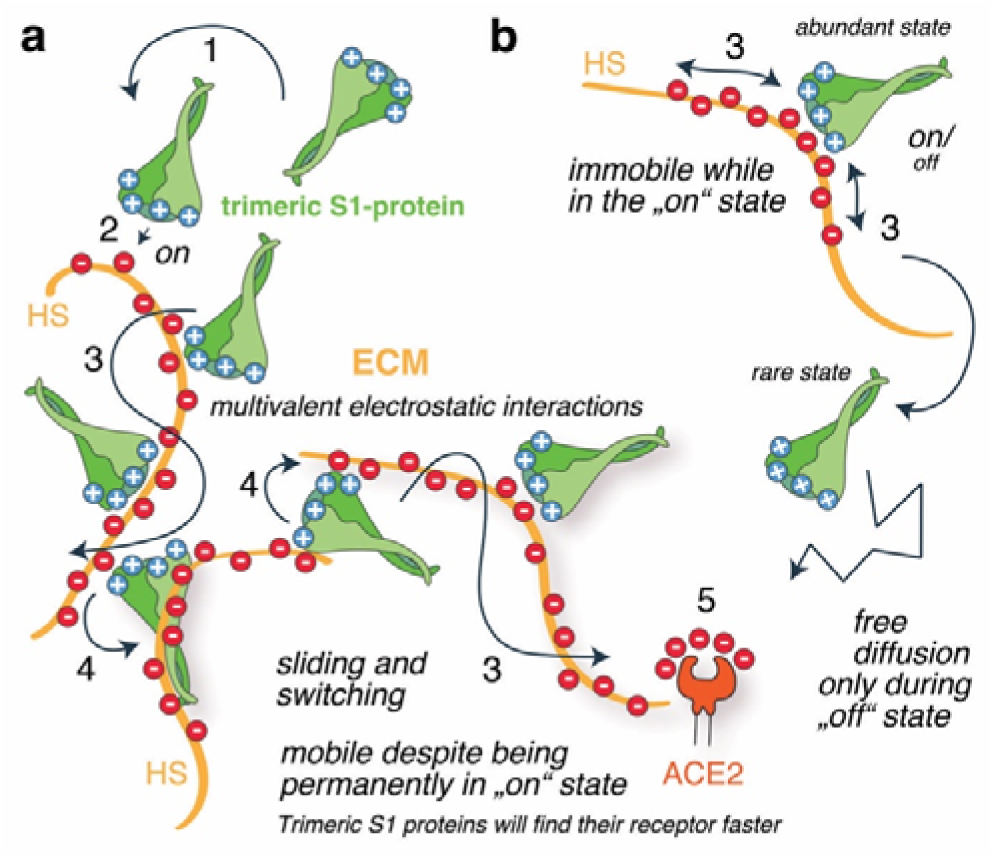
Schematic summary of optimized multivalent and exchangeable S1-protein/HS interactions. **a)** Viral S1-protein interactions with cells involve the following steps: Multivalent attraction (electrostatic steering, 1), precise docking (2), the ability to slide along the polyelectrolyte (3), and the ability to effectively switch between HS to rapidly find ACE2 receptors (4). We show here that SARS-CoV-2 VOCs have optimized the multivalent and electrostatic nature of the trimeric S1-protein/HS interaction so that it allows the protein to partially unbind and rebind the HS chain, but with little tendency to completely unbind the HS, as this would require all individual bonds to be “off” at the same time. **b)** For classical on/off binders, such multivalent association is therefore expected to impair their movement within the ECM, as it reduces their chance to revert into the “off” state required for diffusion and ACE2 binding (5). In this work, we show that the optimized multivalency and dynamic nature of electrostatic protein/HS interactions allows the trimeric S-proteins to relocate within the ECM while remaining in the on-state (Representing the mode depicted in a), likely facilitated by their ability to simultaneously bind at least two HS chains and to directly switch between them (4). We note that SARS-CoV-2 was free to evolve this potential because the properties of the HS binding partner, as well as pH and ion concentrations, remained fixed due to functional constraints. Red: negatively charged sulfates, blue: positively charged amino acids.

This evolution raises the question of whether the reduced *k_off_* of the S-protein/HS interaction conflicts with the requirement for viral movement at the cell surface and ACE2 binding. From the viral evolutionary perspective, an ideal situation would be enhanced binding near the cell surface, combined with an enhanced ability to migrate along and/or between HS chains to locate ACE2 receptors faster. Therefore, the virus ideally evolves its S-protein surface to optimize the initial binding strength (affinity) and the half-life of the interaction such that it remains on the cell surface, and at the same time the ability to migrate along and between HS chains by keeping the interaction dynamic enough. We show that, counterintuitively, these are not conflicting modes, since the increased positive charge of trimeric VOC S-proteins also enhances their ability to switch directly between heparin chains (Figure 7). This important aspect of dynamic HS interactions is fundamentally different between the RBDs and the trimeric S-proteins and has apparently undergone evolutionary optimization. In contrast to the RBD and the trimeric S-proteins of the Wuhan variant, which show similar rates of (low) detachment from the heparin film, regardless of the presence or absence of soluble acceptors, the trimeric Omicron S-protein – in contrast to the RBD – desorbs much faster in the presence of soluble acceptors. We explain this by the multivalency of the S-protein/heparin interaction possibly combined with conformational switching of the trimeric S-protein [56]. Indeed, the multivalent effect posits that trimeric S-protein/HS interactions consist of the sum of many binding modes and sites that are linked and that can partially – but not completely – unbind in order to engage a new interaction. This requires that there must be at least one of several possible interactions between the two molecules at any time, and that the situation is dynamic both in binding terms and conformational terms. *In vivo*, this mechanism could translate into faster mucosal diffusion of the VOCs within the upper airway to the ACE2 target on the epithelium to facilitate efficient infection in the upper airway epithelial cells [11], whereas the delayed ACE2 association of the Wuhan variant may have facilitated subsequent passive transport by diffusion or fluid and mucus flow into the lower airway [47]. Indeed, trimeric S-protein desorption of the Omicron VOC is further enhanced when the negative charge of the soluble acceptor molecule (PPS) is increased. This result indicates that the interaction between trimeric S-proteins and the HS is mainly charge-based and non-specific, i.e. predominantly electrostatic in nature. This raises the interesting possibility that viral surfing may be directed from sites of low HS charge to sites where HS sulfation is increased, and that different extents of lung HS biosynthesis and sulfation may influence individual susceptibility to SARS-CoV-2 infection. Indeed, an important recent publication describes that the composition of human lung HS varies widely among individuals depending on sex, age, and health status, and that compositional differences among samples affected chemokine binding affinities to varying degrees [57]. It is therefore possible that the binding of the virus to the glycocalyx and its movement through the ECM varies between individuals, which may be a factor contributing to the observed differences in disease susceptibility and severity between patients. We also note that the evolutionary optimization described here, and its possible consequences *in vivo*, may not be restricted to SARS-CoV-2 infection but may also apply to other HS-binding virus families.

In a final set of experiments, we exploited the evolutionary optimized electrostatic S-protein/HS interactions as a potential druggable target. It has been shown previously that heparin [11, 29, 33, 43] and PPS inhibit viral interaction with cellular HS and infection to a similar extent [33, 46]: The IC_50_ of PPS against the RBD of the Wuhan variant was determined to be ∼35 nM and the IC_50_ of heparin against the RBD was determined to be 56 nM [46]. Treatment of African green monkey Vero kidney epithelial cells (Vero E6 cells) with heparinases to reduce cell surface HS also reduced the percentage of infected cells by ∼5-fold, demonstrating the importance of initial HS contact for SARS-CoV-2 infection [11, 29]. Extending these previous studies, we asked whether the accumulation of positive charge during S-protein evolution altered the ability of heparin or PPS to inhibit cell surface binding and infection. PPS has previously been shown to inhibit Wuhan variant invasion in experiments involving all three forms of addition; PPS added to cells, to virus, and when added to both before mixing [33]. In these experiments, the inhibitory capacity of PPS was equal to or better than that of heparin [33]. We confirmed this finding under all three conditions and also found that heparin inhibited the infection of Vero cells by SARS-CoV-2 Wuhan and Omicron variants to a similar extent. However, we also found that cell pre-treatment and cell and virus pre-treatment with PPS inhibited infection by the Omicron variant significantly more than infection by the Wuhan variant, and cell and virus pre-treatment almost completely inhibited the Omicron variant. This indicates that the accumulation of additional positive charge on the Omicron S-protein has increased its susceptibility to highly negatively charged soluble inhibitors, which are likely to act on three mechanisms of SARS-CoV-2 infection: allosteric or charge-based inhibition of virus binding to ACE2, enhanced competitive inhibition with binding to host HS coreceptors and, possibly, its desorption from cellular HS of lesser charge and prevention of spike cleavage at polybasic sites [58]. Together with the fact that PPS has about 10 times lower anticoagulant potency than heparin and no anti-factor Xa activity [44, 45], the administration of PPS, even at high doses, may therefore allow its use as a preventive measure, i.e. in nasal sprays to inhibit viral attachment to the glycocalyx, to increase viral desorption and to decrease cellular rebinding, in particular of the Omicron VOCs and future variants.

## Materials and Methods

### Molecular docking analysis

#### Protein Preparation

The structures of RDB variants (residues 333-526) for docking calculations were prepared using the *“Protein preparation”* wizard of Schrödinger-Maestro (release 2023-1), starting from the corresponding X-ray structures. For all the three variants, the active open prefusion conformation (RBD-accessible) structure in the complex with the ACE2 receptor was retrieved from the PDB database: the X-ray structures were selected for the wild-type (WT, PDB ID: 6M0J [7]) and the Omicron variant (BA 1.1.259, PDB ID: 7WBP [59]) and the cryo-electron microscopy structure (PDB ID: 7V8B) for the Delta variant (B1.617.2 subline). We then added the N-glycosylation at residue N343 using the WT RBD-accessible structure of the fully glycosylated homotrimeric SARS-CoV-2 spike head (PDB ID: 6vsb_1_1_1, chain A) as a template. The structure was obtained from the CHARMM-GUI Covid-19 protein archive and it is based on the cryo-EM structure of the RBD-accessible state (PDB: 6VSB). The glycan attached to the N343 is an octasaccharide with the formula [GlcNA(α1-6)Fuc](β1-4)GlcNAc(β1-4)Man[(β1-3)Man(α1-6)GlcNAc][(β1-6)Man(α1-6)GlcNAc], which is the most abundant sequence found for all three variants. After removal of all the crystallographic water, the hydrogen atoms were added and disulfide bonds between cysteines were formed. For the protein residues, pKa values were calculated using the PROPKA method at pH 7.4. Hydrogen bonds were then optimized using the exhaustive sampling option. Finally, the structures were relaxed by using a restrained minimization with convergence on heavy atoms to an RMSD (root-mean-square deviation) of 0.30 Å and the OLPS_2005 force field.

#### Ligand Preparation

The 3D structures of the ligands were taken from our previously published work. Specifically, the hexasaccharide α-l-IdoA2S(1→4)α-d-GlcNS,6S(1→4)β-d-GlcA(1→4)α-d-GlcNS,6S(1→4)α-l-IdoA2Sα-d-GlcNS,6S(1)-OMe ligand was constructed considering the ^4^C_1_ chair conformation of glucosamine and glucuronic acid residues and the ^1^C_4_ chair conformation for IdoA residues. The glycosidic dihedral angles, in order from non-reducing to reducing residue, were set as follows: *ψ*_l_/*ϕ*_l_ = 40°/-14° *ψ*_2_/*ϕ*_2_=-40°/-30° *ψ*_3_/*ϕ*_3_=60°/30° *ψ*_4_/*ϕ*_4_ =-33°/-39° *ψ*_5_/*ϕ*_5_=41/14°. The conformation of the sulfate groups of GlcNS6S and IdoA2S was set in accordance with the experimental heparin structure (PDB ID:1HPN). The ligand has a net charge of -11. The linear hexasaccharide PPS was built using the polysulfate xylose unit in ^4^C_1_ chair conformation: this chair conformation is characterized by sulfate groups in the axial position, which allows minimizing the strong electrostatic repulsion between these negatively charged groups. While the first five monosaccharide residues (Xyl-1 to Xyl-5) are 3,4- and 2,3-disulfo xylose units, the last one (Xly-6) has an additional O-sulfation at position 4, resulting in a total charge of -13.

#### Docking Setup

Automated docking calculations were performed by using Glide (version 80012, OPLS_2005 force field) in Standard Precision (SP) mode. The proteins were considered as rigid bodies, while the ligands were free to vary their conformation and orientation within the binding site. To avoid inaccurate conformational sampling, the ring conformation and nitrogen inversion options were not applied and all the f and y torsions around all glycosidic bonds were kept fixed. We increased the number of poses generated in the initial stage of docking to 10,000 and saved 5,000 poses for the energy minimization step. No other changes were made to the default settings. The receptor grids were generated on the prepared proteins using the OPLS_2005 force field with the outer cubic box of 55 Å and the inner box of 25 Å centered on the barycenter of protein residues R346, N354, R355, K356, R357, R466. For each ligand, the Glide scoring function was used to select the top 10 poses after a post-minimization step. The docking poses were also re-scored using Molecular Mechanics Generalized Born Surface Area (MM-GBSA) binding energy calculations with Prime software 13 (Prime MM-GBSA version 3.0) and the VSGB2.0 14 implicit water model with the OPLS_2005 force field (Table S1). The contact analysis was performed using the *Interaction fingerprint* tool available on Schrödinger [60]. The default criteria for hydrogen bond formation are: maximum distance between H-bond donor (HBD) and acceptor (HBA) is 2.8 Å, minimum HBD and HBA angles are 120° and 90° respectively. In case of salt bridges, the maximum distance required for positive interaction between charged groups is 5 Å.

### QCM-D

#### Reagents

The following S1-RBDs were used in this study: RBD of the first ermerged variant (*Wuhan*, Sinobiological 40592-V08H), of the B.1.167.2 variant (*Delta*, Sinobiological 40592-V08H90), and of the B.1.1.529 variant (*Omicron*, Sinobiological 40592-V08H121). S1+S2 homo-trimers of the Wuhan variant (Sinobiological 40589-V08H8) and the Omicron variant (Sinobiological 40589-V08H26) were also analyzed. Size exclusion chromatography (SEC) was always used to confirm the trimeric state of the S-proteins prior to testing by QCM-D. Heparin and HS were obtained from porcine intestinal mucosa (Sigma-Aldrich and Celsus Laboratories, Cincinnati, OH, USA); HS carrying 1.4 sulfates/disaccharide was further purified in the laboratory of Hughes Lortat-Jacob (Institut de Biologie Structurale, Université Grenoble Alpes, Grenoble, France) [61]. The average mass of all the heparin and HS isolates, when anchored to the surface, was estimated to be 9 kDa (∼ 18 disaccharide units) by QCM-D analysis [62]. Pentosan polysulfate (PPS L26, Mw = 4890 g/mol) was kindly provided by Bene Pharma and used at the indicated concentrations. A single molecule of PPS has been described to bind four to five S1 RBDs [33]. PPS has poor anticoagulant activity and has been approved for the treatment of bladder pain and discomfort in interstitial cystitis (ema.europa.eu/en/medicines/human/EPAR/elmiron).

#### Synthesis of biotinylated heparin

Biotinylation of heparin was performed as previously described [40], by adapting a previously reported procedure [63]. Briefly, heparin (4 mM) was dissolved in 100 mM acetate buffer (prepared from glacial acetic acid (Carl Roth, Karlsruhe, Germany) and sodium acetate (Sigma-Aldrich) at pH 4.5) containing aniline (100 mM, Sigma-Aldrich). Biotin-PEG_3_-oxyamine (3.4 mM, Conju-Probe, San Diego, USA) was added to the heparin solution and allowed to react for 48 h at 37 °C. The final product was dialyzed against water for 48 h by using a dialysis membrane with a 3.5 kDa cutoff. The final solution was then lyophilized and stored at -20 °C. For further use, the conjugates were diluted to the desired concentrations in buffer. The resulting biotinylated heparin was characterized by biotin-streptavidin binding assays using QCM-D. Low-sulfated and high-sulfated HS was treated in the same manner.

#### Preparation of small unilamellar vesicles (SUVs)

SUVs were prepared by adapting reported procedures [64, 65], as described previously [40]. Briefly, a lipid mixture consisting of 1 mg/mL 1,2-dioleoyl-sn-glycero3-phosphocholine (DOPC, Avanti Polar Lipids) and 5 mol% of 1,2-dioleoyl-sn-glycero-3-phosphoethanolamine-N-(cap biotinyl) (DOPE-biotin, Avanti Polar Lipids) was prepared in chloroform in a glass vial. The solvent was then evaporated with a weak stream of nitrogen while the vial was rotated to obtain a homogeneous lipid film. The residual solvent was removed under vacuum for 1 h. The dried film was then rehydrated in ultrapure water to a final concentration of 1 mg/mL and vortexed to ensure the complete solubilization of the lipids. The lipids were sonicated for approximately 15 min until the opaque solution became clear. The resulting SUVs were stored in the refrigerator and used within 2 weeks. For the FRAP experiments, lipid mixtures of DOPC and DOPE-biotin with DiD (Sigma Aldrich) (molar ratio 94.9 : 5.0 : 0.1) were used to form the SUVs.

#### QCM-D measurements

QCM-D measurements were performed with a QSense Analyser (Biolin Scientific, Gothenburg, Sweden) and SiO_2_-coated sensors (QSX303, Biolin Scientific). Measurements were performed at a working temperature of 23 °C with four parallel flow chambers and a peristaltic pump (Ismatec, Grevenbroich, Germany) with a flow rate of 20 µL per min. In some experiments, a multi-channel syringe pump was instead used to control the flow, at a rate of 20 μL/min. The normalized frequency shifts Δ*F*, and the dissipation shifts Δ*D*, were measured at six overtones (*i* = 3, 5, 7, 9, 11, 13). The fifth overtone (*i* = 5) was presented throughout except for QCM-D binding data obtained at different S-trimer concentrations, where the seventh overtone was used; all other overtones gave qualitatively similar results. QCM-D sensors were first cleaned by immersion in a 2 wt% sodium dodecyl sulfate solution for 30 min, followed by rinsing with ultrapure water. The sensors were then dried under a nitrogen stream and activated by a 10 min treatment with a UV/ozone cleaner (Ossila, Sheffield, UK). For the formation of supported lipid bilayers (SLBs), after obtaining a stable baseline, freshly prepared SUVs were diluted to a concentration of 0.1 mg/mL in buffer solution (wash buffer A, 50 mM Tris, 100 mM NaCl (Sigma Aldrich) at pH 7.4) containing 10 mM of CaCl_2_ just before use and flushed into the chambers. The quality of the SLBs was monitored *in situ* to ensure that high-quality SLBs were formed, corresponding to equilibrium values of Δ*F* = -24 ± 1 Hz and Δ*D* < 0.5 ppm. A solution of streptavidin (SAv; 150 nM) was then passed over the SLBs, followed by the addition of biotinylated heparin (10 μg/mL, except for QCM-D binding data obtained at different S-trimer concentrations, where 13 μg/mL was used). Each sample solution was washed over the QCM-D sensor until the signals equilibrated and then rinsed with wash buffer A (see above). Prior to the addition of SARS-CoV-2 S1 RBD or trimer solutions 200nM, 5µg/ml), the flow rate was reduced to 20 µL/min. For titrations, increasing concentrations of proteins were used as indicated, as well as for the heparin and PPS solutions. Data were acquired with QSoft401 version 2.7.3.883, and analyzed in Origin.

#### FRAP measurements

For the FRAP experiments, SLBs were deposited in 18-well glass bottom microslides (Ibidi, Gräfelfing, Germany). Prior to SLB formation, 150 μL of aqueous 2 M sodium hydroxide solution was added to the glass substrate for 1 h to form a hydrophilic surface. The wells were then rinsed three times with ultrapure water and three times with buffer (50 mM Tris, 100 mM NaCl, pH 7.4) containing 10 mM CaCl_2_. A solution of 0.2 mg/mL SUVs in buffer containing 10 mM CaCl_2_ was then added for 30 min at room temperature. An SLB was then formed by the process of binding, rupture and spreading of the SUVs on the glass substrate. Excess lipids were removed from the well by five washes with 100 μL buffer. After the SLB formation, 50 μL buffer was left in the well plate in order to preserve the SLBs. A solution of Cy3-streptavidin (Cytiva, final concentration 1.5 μM) and biotinylated-heparin (final concentration 100 μg/mL) was then added to the well plates for 30 min. Finally, solutions of Omicron S trimer (final S-protein trimer concentrations were set 2-fold the concentrations of half-maximal QCM-D responses, Supplementary Figure 6a) were added for another 30 min, and excesS-proteins were washed off five times with 100 μL buffer. Using a confocal microscope, a circular spot of approximately 25 µm in diameter was bleached with a laser at 552 nm and 638 nm (100% intensity), and the fluorescence intensity in the bleached regions was monitored. For both the SLB and the SAv, the FRAP protocol consisted of 3 imaging loops (0.7 s intervals) before bleaching, 10 loops during bleaching (0.7 s intervals), and 20 loops during recovery (10 at 0.73 s intervals and 10 at 10 s intervals). All FRAP measurements were performed with a Leica SP8 confocal laser scanning microscope through a 63× water objective. The Cy3 dye was excited with a 552 nm laser, and the emission was detected at 560 – 630 nm. All images were analyzed using the Leica Application Suite X (LAS X) software version 3.7.1.21655. Data were plotted using Origin Lab version 2022b (9.9.5.167).

### SARS-CoV-2 infection

Vero cells were plated at 2.5×10^5^ cells/well in 24-well plates in Eagle’s Minimum Essential Medium (EMEM) supplemented with 10% fetal bovine serum (complete medium). The advantage of these cells is that they are well characterized and do not secrete α- or β-interferon when infected by the virus. After 24 h, the cells were infected with SARS-CoV-2 isolate (D614G) (GISAID accession ID: EPI_ISL_413489) or Omicron BA.1 isolate (GISAID accession ID: EPI_ISL_12221510), which was previously titered by the plaque assay in Vero cells [66, 67]. Heparin and PPS were added according to three different protocols. i. Cells were incubated with 100 µg/ml of either heparin or PPS in 250 µl of complete medium for 30 min and then 50 µl of supernatant containing 50 plaque-forming units (PFU) was added to Vero cells (cell pretreatment). ii. 50 PFU of SARS-CoV-2 in 50 µl of complete medium were incubated with compounds (100 µg/ml, a heparin concentration within the range experienced during anticoagulation therapy) for 30 min at 37 °C and then added to Vero cells in a final volume of 300 µl (virus pre-treatment). iii. 50 PFU of SARS-CoV-2 in 50 µl of complete medium were incubated with compounds (100 µg/ml) for 30 min at 37 °C and cells were also incubated with compounds (100 µg/ml) in 250 µl of complete medium for 30 min at 37 °C. After this incubation period, the virus-containing supernatant was added to the cells (cell + virus pre-treatment). In all three protocols, the incubation was extended for 1 h at 37°C. The supernatants were then discarded, and 500 µl of 1% methylcellulose overlay dissolved in medium containing 1% of fetal bovine serum was added to each well. After 3 days, cells were fixed with 6% (v/v) formaldehyde phosphate-buffered saline and stained with 1% (w/v) crystal violet (Sigma-Aldrich) in 70% (v/v) ethanol (Sigma-Aldrich). Plaques were counted under a stereoscopic microscope (SMZ-1500, Nikon). Prism GraphPad v. 9.0 software (www.graphpad.com) was used for the statistical analyses.

## Acknowledgements

SFB1348A08 funding to K.G., BBSRC Research Grant BB/X007278/1 to R.P.R. We thank H. Lortat Jacob (Institut de Biologie Structurale, Université Grenoble Alpes, Grenoble, France) for providing the HS.

## Notes

### Competing Interest Statement

The authors have declared no competing interest.

