## Supplemental Information for "Evolution of SARS-CoV-2 spike trimers towards optimized heparan sulfate cross-linking and inter-chain mobility"

**Supplementary Figures:**

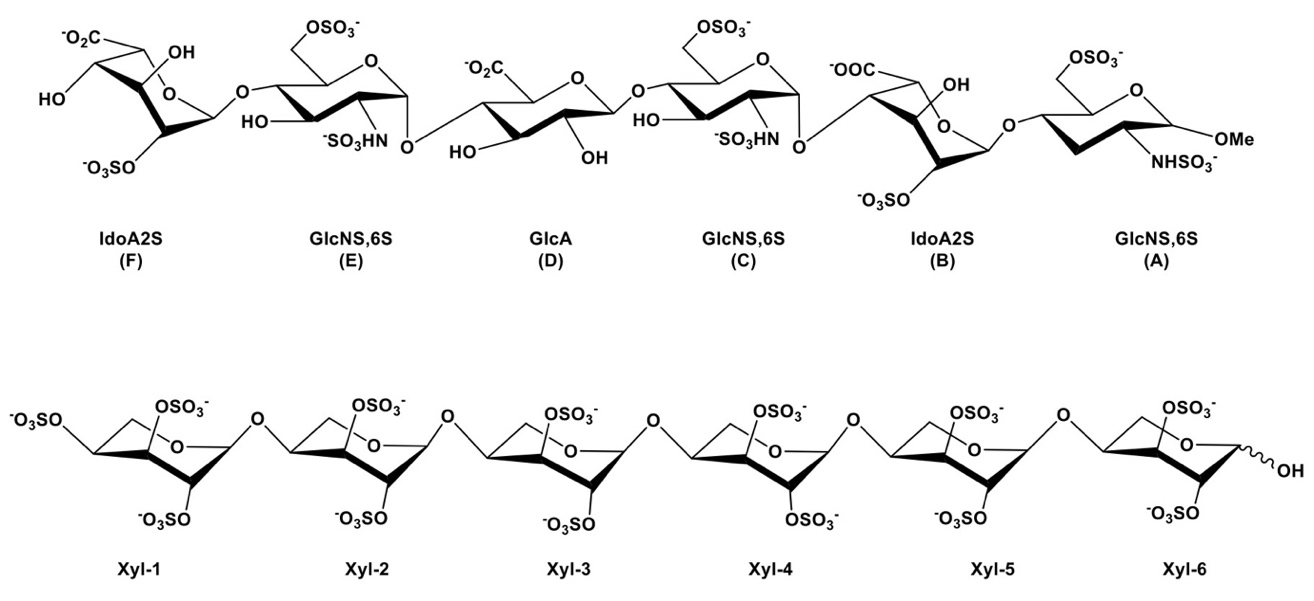

**Supplementary Figure 1:** Structure of IAGAIA (top) and PPS (bottom) ligands used for docking experiments with RDB protein VOCs.

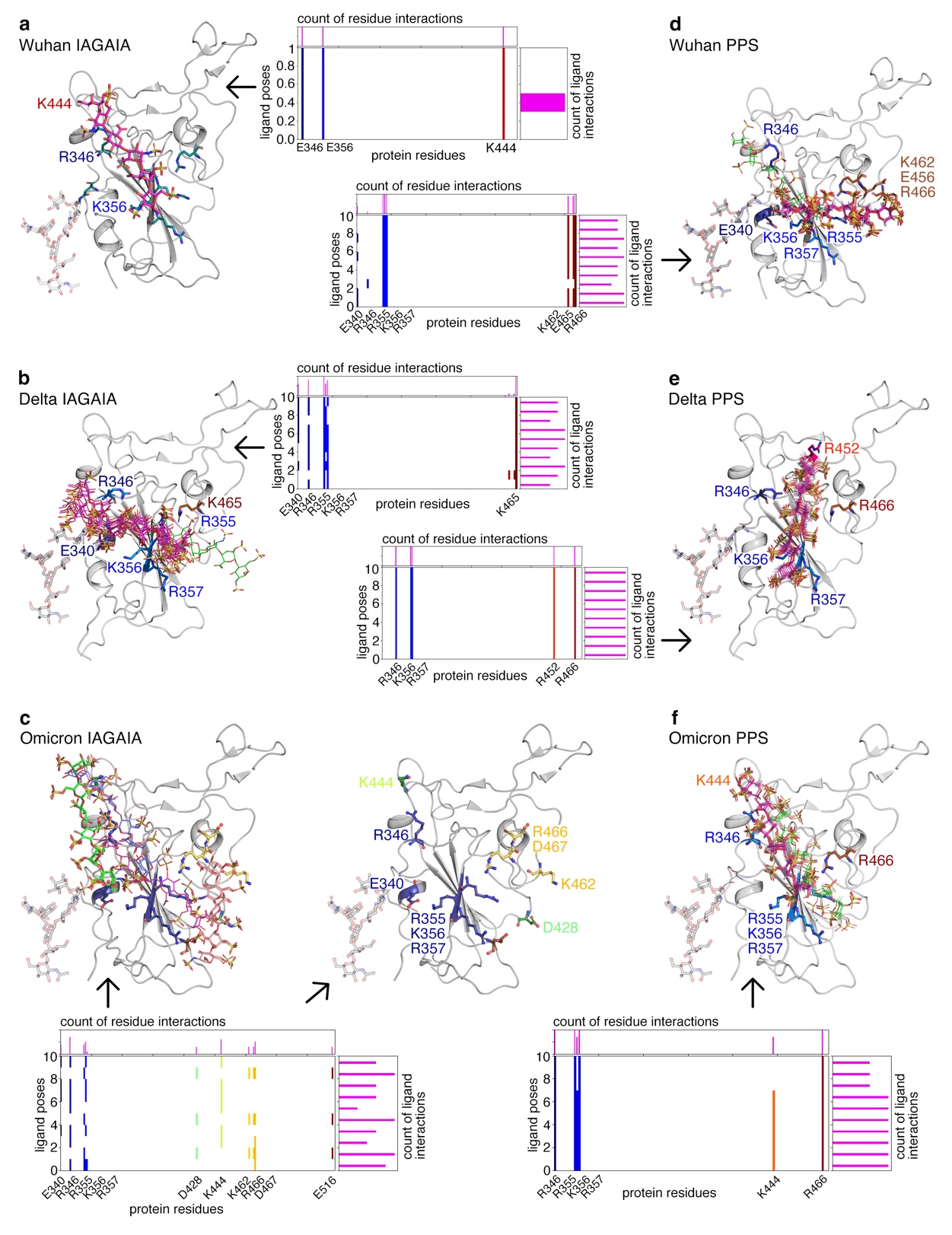

**Supplementary Figure 2: Analysis of the docking pose interactions of IAGAIA (a,c,e) and PPS (b,d,f) in the Wuhan, Delta, and Omicron RBD models. Key residues are labelled in the figure and for each docking pose the charged residues making interactions with the ligand are shown in the graph below.** Moving from IAGAIA to PPS the ligand is prone to exhibit a smaller number of binding modes due to the more focused interaction with basic residues exposed in the basic channel located in the binding site.

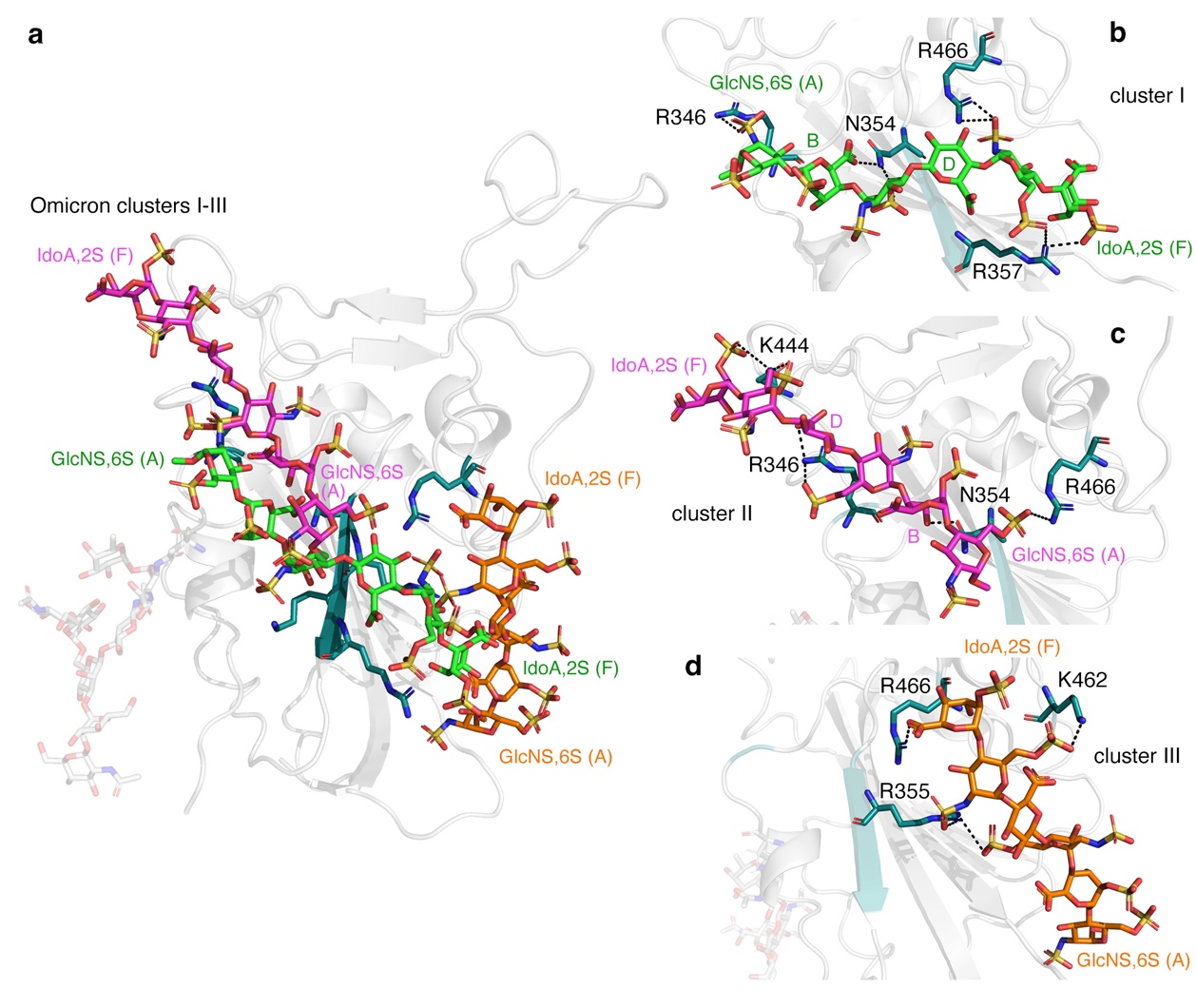

**Supplementary Figure 3: a)** IAGAIA and Omicron RBD cluster I (green), II (magenta), and III (orange). **b-d)** Best se of cluster I, II and III. The RBD region is represented by a grey cartoon, while IAGAIA represented red, blue and yellow sticks indicating oxygen, nitrogen and sulfur atoms, respectively. The key interacting residues of the pocket are labelled and depicted with similar color codes (deep teal for carbon, red for oxygen and blue for nitrogen), while H bond and salt bridges are represented as black dashed lines.

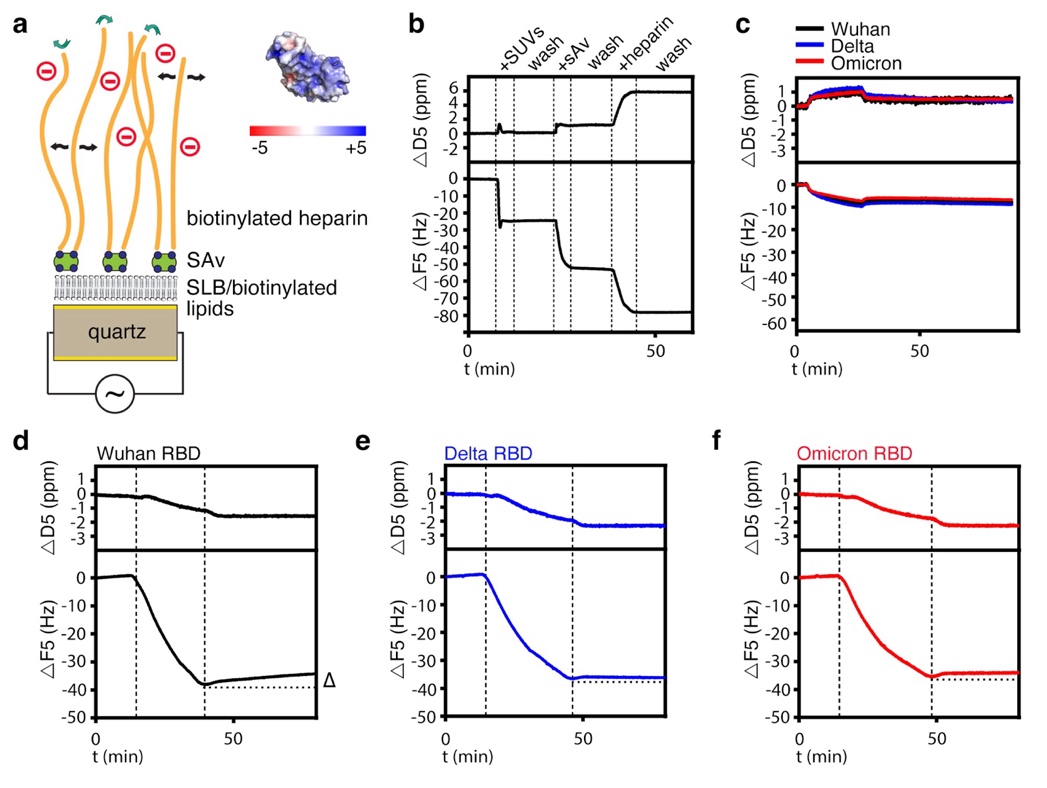

**Supplementary Figure 4:** **Quartz crystal microbalance with dissipation monitoring (QCM-D).** **a)** The core of the QCM technology is a gold-coated oscillating quartz crystal sensor disk with a resonance frequency related to the mass of the disk. This allows the real-time detection of nanoscale mass changes on the sensor surface by monitoring changes in the resonance frequency (Δ*F*). Interaction surfaces containing biotinylated (blue) heparin (orange) linked via steptavidin (green) to fluid supported lipid bilayers (SLBs, black) were generated as a proxy for cell-surface-bound HS. Like cell surface HS, SLB-linked highly negatively charged heparin can freely rotate (green curved arrows) and potentially move laterally (black arrows) on the sensor surface. Adsorption of positively charged molecules on the heparin-covered surface decreases *F*, and mass loss during washing increases *F*. QCM-D measures an additional parameter, the change in energy dissipation *D*, which is particularly useful for studying viscoelastic properties of the layer. An increased Δ*D* during protein binding to the functionalized surface correlates with a softer layer, and a decreased Δ*D* would indicate layer stiffening, for example by cross-linking of heparin chains by the bound molecules. **b)** Preparation and validation of a heparin/HS cell surface matrix model for QCM-D. Representative QCM-D data displaying the observed frequency (Δ*F*) and dissipation (Δ*D*) shifts during assembly of the SLB, the SAv monolayer and the film of end-attached heparin on the sensor’s silica surface. Start and duration of sample incubations are indicated by dashed vertical lines and a label on top of the graph. At all other times, the surface was exposed to wash buffer (10 mM HEPES pH 7.4, 150 mM NaCl). The formation of the heparin/HS model matrix on the QCM sensor was always followed in real-time prior to the protein incubation assays shown in Figures 3-5 and Supplementary Figures 2-4 to validate the surface functionalization. **c)** A control addition of RBDs to a surface without terminally attached heparin shows much smaller responses than on heparin, confirming that RBD binding to heparin is largely specific. **d-e)** Representative QCM-D data (shown analogous to Fig. 3a) for model matrix formation with heparin and subsequent RBD interaction to better discriminate VOC unbinding responses. The start and duration of sample incubations are indicated by dashed vertical lines and a label at the top of the graph. At all other times, the surface was exposed to wash buffer. Note the overall similarity in RBD binding to heparin, but the different stability of the interaction when washed in buffer (the dotted horizontal line represents -Δ*F* at the start of the buffer wash).

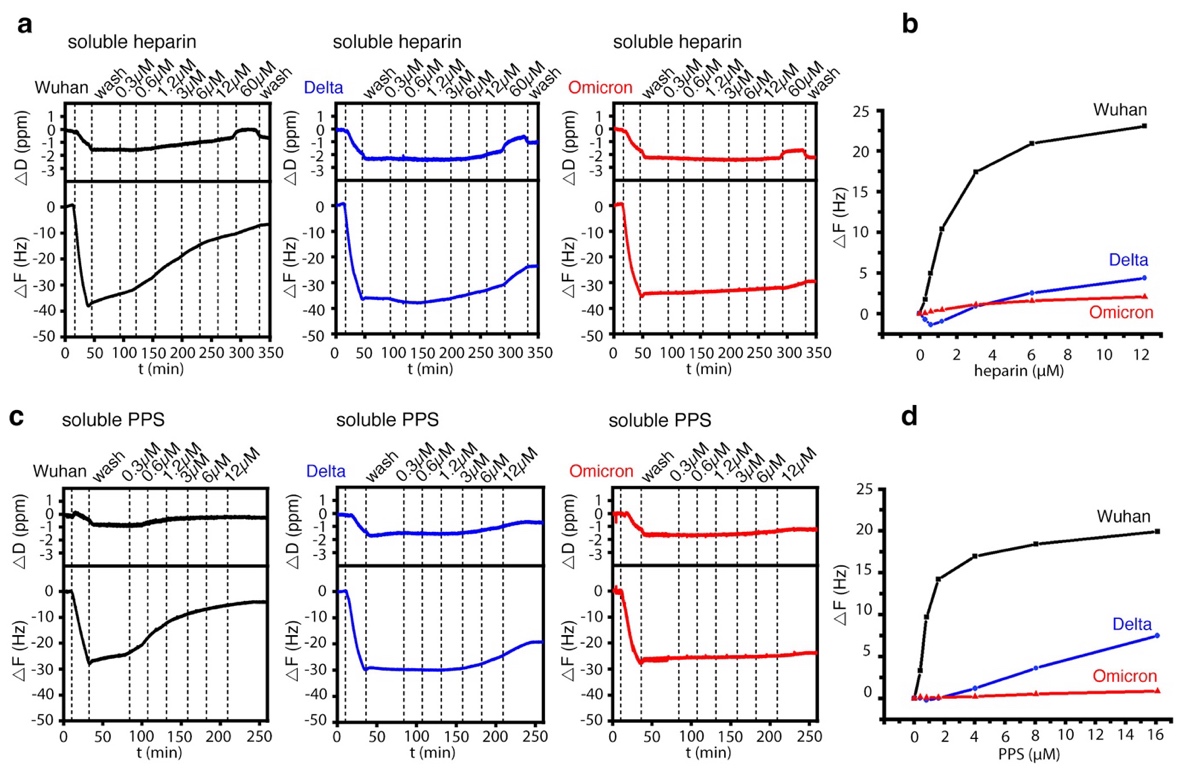

**Supplementary Figure 5: Soluble heparin and PPS increase the dissociation of the Wuhan and Delta RBDs, but not that of the Omicron RBD. a)** Individual graphs of data shown in Figure 4a. The dissociation of the Wuhan S-protein RBD from the sensor surface increases with increasing heparin concentration in the washing buffer, and the Delta S-protein RBD also dissociates at higher heparin concentrations. The Omicron RBD does not dissociate from the heparin-functionalized sensor surface. **b)** Relative heparin elution strength for the Wuhan, Delta and Omicron RBDs from the sensor surface. The starting point of the curves is represented by the -ΔF value at the end of the wash step. **c)** As shown in Figure 4b, increasing the negative charge of soluble PPS in the wash buffer increases the dissociation of the Wuhan and Delta RBDs, but not of the Omicron RBD. **d)** Relative PPS elution strength for the Wuhan, Delta and Omicron RBDs from the sensor surface.

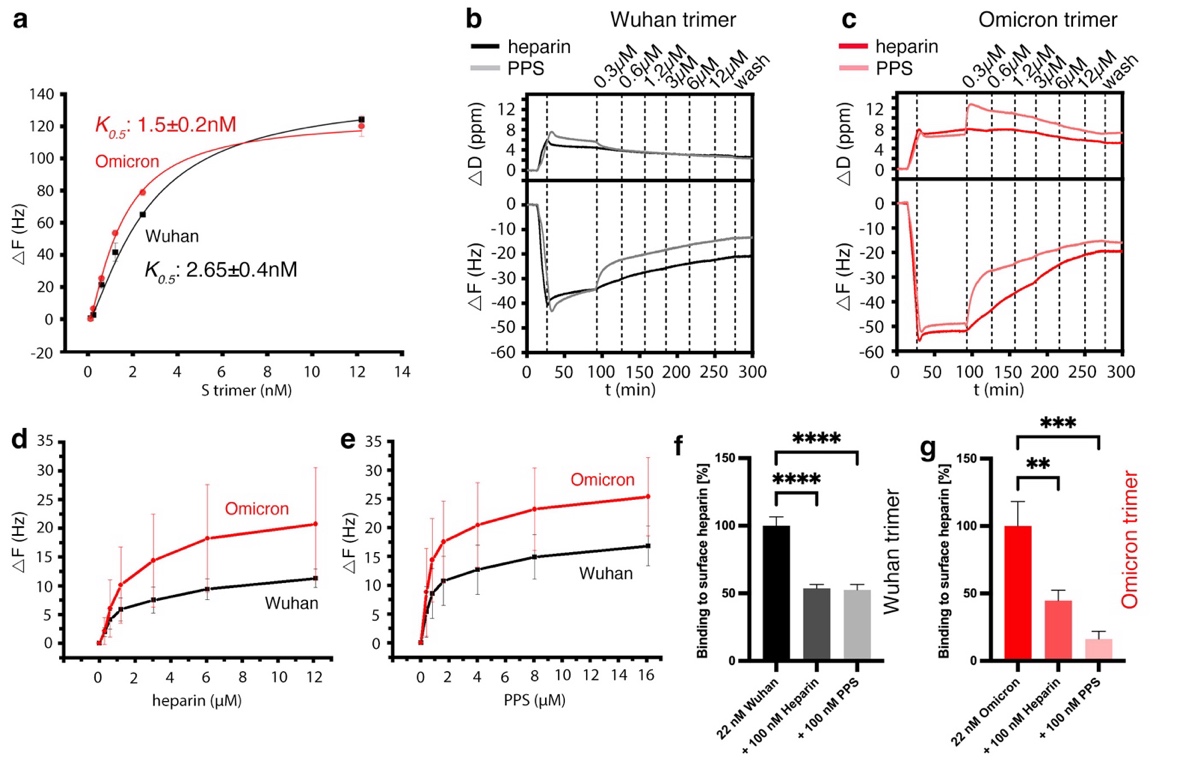

**Supplementary Figure 6: Strongly increased Omicron S-trimer switching to the soluble PPS acceptor indicates charge-driven direct switching of the protein.** **a)** Hill fit of QCM-D binding data obtained at different S-trimer concentrations. The concentration of half-maximal binding *K*_0.5_ of the Wuhan S-trimer exceeded that of the Omicron variant by a factor of approximately 2. **b)** Low amounts of soluble heparin and PPS readily desorb Wuhan S-trimers from the heparin-functionalized sensor surface, with PPS being more effective (note the steep slope immediately after the addition of soluble PPS, gray line, asterisk). **c)** The same low amounts of soluble heparin and PPS desorb Omicron S-trimers much more strongly from the heparin-functionalized sensor surface, with PPS again being most effective. Note that he striking increase in trimeric Omicron S-protein desorption was accompanied by a concomitant increase in *D*. One possibility to explain this result is a (non-specific) association of PPS with the sensor surface. However, this possibility is unlikely because of the strong repulsion between the equally charged PPS and the immobilized heparin. It is also unlikely because higher amounts of PPS in the wash buffer do not further increase *D*. Therefore, we suggest that the observed increase in *D* may be due to the rapid de-crosslinking of the heparin film by the trimeric S-protein as a prerequisite for their subsequent switch to the more negatively charged PPS and effective release of the protein/PPS complex into the solution phase. **d)** Relative heparin elution strength for the Wuhan and Omicron S-protein trimers from the sensor surface. The starting point of the curves is represented by the -ΔF value at the end of the wash step. **e)** Relative PPS elution strength for the Wuhan and Omicron S-protein trimers from the sensor surface. **f)** Relative inhibition of Wuhan S-protein trimer association to the sensor surface upon protein preincubation with or without 100nM soluble heparin or PPS. One-way ANOVA, Dunnet’s multiple comparison test, n=3. ****: p<0.0001. **g)** Relative inhibition of Omicron S-protein trimer association to the sensor surface in the absence or presence of 100nM heparin or PPS. One-way ANOVA, Dunnet’s multiple comparison test, n=1 for the Omicron control, n=2 for the PPS-treated protein and n=3 for the heparin-treated probe.

**Supplementary Tables:**

| IAGAIA | | | |
| --- | --- | --- | --- |
| RDB | Pose | Docking Score  (kcal/mol) | MMGBSA  (kcal/mol) |
| **WT (6M0J)** | 1 | -1.51 | -17.67 |
| **Delta (7V8B)** | 1 | -3.95 | -35.27 |
|  | 2 | -3.58 | -34.40 |
|  | 3 | -2.76 | -28.71 |
|  | 4 | -2.72 | -31.74 |
|  | 5 | -2.52 | -27.66 |
|  | 6 | -2.52 | -34.80 |
|  | 7 | -1.91 | -18.26 |
|  | 8 | -1.74 | -18.41 |
|  | 9 | -1.67 | -15.61 |
|  | 10 | -1.03 | -13.79 |
| **Omicron (7WBP)** | 1 | -5.09 | -37.40 |
|  | 2 | -4.92 | -35.35 |
|  | 3 | -4.80 | -31.02 |
|  | 4 | -4.63 | -42.44 |
|  | 5 | -4.53 | -33.34 |
|  | 6 | -4.41 | -35.70 |
|  | 7 | -4.08 | -31.69 |
|  | 8 | -3.97 | -34.41 |
|  | 9 | -3.77 | -24.99 |
|  | 10 | -3.77 | -29.84 |

**Supplementary Table 1:** IAGAIA re-scored MMGBSA values and docking scores for the poses obtained for the Wuhan strain RBD and Delta and Omicron RBDs.

| PPS | | | | |
| --- | --- | --- | --- | --- |
| RDB | Pose | Docking Score  (kcal/mol) | | MMGBSA  (kcal/mol) |
| **WT**  **(6M0J)** | 1 | -4.74 | | -19.21 |
|  | 2 | -4.22 | | -26.41 |
|  | 3 | -4.15 | | -24.43 |
|  | 4 | -3.85 | | -25.16 |
|  | 5 | -3.85 | | -27.08 |
|  | 6 | -3.82 | | -24.91 |
|  | 7 | -3.78 | | -20.75 |
|  | 8 | -3.74 | | -27.33 |
|  | 9 | -3.72 | | -25.31 |
|  | 10 | -3.72 | | -21.72 |
| **Delta**  **(7V8B)** | 1 | -5.13 | | -29.08 |
|  | 2 | -4.85 | | -24.63 |
|  | 3 | -4.70 | | -23.72 |
|  | 4 | -4.50 | | -31.08 |
|  | 5 | -4.49 | | -27.32 |
|  | 6 | -4.47 | | -15.36 |
|  | 7 | -4.47 | | -28.92 |
|  | 8 | -4.29 | | -28.94 |
|  | 9 | -4.24 | | -30.62 |
|  | 10 | -4.11 | | -31.04 |
| **Omicron**  **(7WBP)** | 1 | -5.76 | | -47.28 |
|  | 2 | -5.55 | | -44.35 |
|  | 3 | -5.19 | | -42.98 |
|  | 4 | -5.16 | | -46.56 |
|  | 5 | -5.09 | | -59.84 |
|  | 6 | -5.07 | | -42.28 |
|  | 7 | -5.02 | | -51.16 |
|  | 8 | -3.67 | | -35.94 |
|  | 9 | -3.22 | | -32.38 |
|  | 10 | -3.15 | | -38.95 |

**Supplementary Table 2:** PPS re-scored MMGBSA values and docking scores for the poses obtained for the Wuhan strain RBD and the Delta and Omicron RBDs .

|  |  | **Protein residue** | **Ligand residue** | **Distance** |
| --- | --- | --- | --- | --- |
| IAGAIA | **Wuhan** | K444-**N**H_3_^+^  K356-**N**H_3_^+^  R346-NH**C**NH_2_ ^+^ | IdoA2S(F)-**C**OO^-^  GlcNS,6S(A)-N-**S**O_3_^-^  GlcNS,6S(E)-3-**O**H | 4.2Å (SB)  3.8Å (SB)  3.9Å (HB) |
|  | **Delta** | R357-NH**C**NH_2_^+^  R357 C_α_**N**H  R346-NH**C**NH_2_^+^  R346-NH**C**NH_2_^+^  N354-C**N**H_2_  N354-C**N**H_2_ | IdoA2S(F)-2-O-**S**O_3_^-^  IdoA2S(F)-3-**O**H  IdoA2S(B)**-C**OO^-^  GlcNS,6S(C)-6-O-**S**O_3_^-^  GlcNS,6S-N-**S**O_3_^-^ (E)  GlcNS,6S-6-O-**S**O_3_^-^ (C) | 3.5Å (SB)  3.1Å (HB)  4.4Å (SB)  4.0 Å (SB)  4.6 Å (SB)  3.6 Å (SB) |
|  | **Omicron Cluster I** | R346-C_α_**N**H  R357-NH**C**NH_2_^+^  N354-C_α_**N**H_2_  N354-C_α_**N**H_2_  R466-HN**C**NH_2_^+^  R357- HN**C**NH_2_^+^ | GlcNS,6S-N-**S**O_3_^-^  IdoA2S-2-O-**S**O_3_^-^  IdoA2S(B)-**C**OO^-^  GlcNS,6S(C)-6-O-**S**O_3_^-^  GlcNS,6S(E)-N-**S**O_3_^-^  GlcNS,6S(E)-6-O-**S**O_3_^-^ | 4.7Å (HB)  5.0Å (SB)  4.5Å (HB)  3.4Å (HB)  4.0 Å (SB)  4.8 Å (HB) |
|  | **Omicron Cluster II** | R466-HN**C**NH_2_^+^  K444-**N**H_3_^+^  K444-**N**H_3_^+^  R346-HN**C**NH_2_^+^  R346-HN**C**NH_2_^+^ | GlcNS,6S(A)-4-O-**S**O_3_^-^IdoA2S(F)-O-2-**S**O_3_^-^  GlcNS,6S(E)-6-O-**S**O_3_^-^  GlcA(D)-**C**OO^-^  GlcNS,6S(C)-6-O-**S**O_3_^-^ | 4.0 Å (SB)  4.0Å (SB)  3.2Å (SB)  3.3Å (SB)  3.8 Å (SB) |
|  | **Omicron Cluster III** | R355-HN**C**NH_2_^+^  R355-HN**C**NH_2_^+^  R466-HN**C**NH_2_^+^  R466-C_α_**N**H  K462-**N**H_3_^+^ | GlcNS,6S(E)-N-**S**O_3_^-^  GlcNS6S-6-O-**S**O_3_^-^  IdoA2S(F)-**C**OO^-^  IdoA2S(F)-3-**O**H  GlcNS,6S(E)-6-O-**S**O_3_^-^ | 4.7Å (HB)  4.0 Å (SB)  4.9Å (HB)  2.9Å (HB)  3.5Å (SB) |
| PPS | **Wuhan Cluster I** | K462**-N**H_3_^+^  K462**-N**H_3_^+^  N354-C**N**H_2_  K356-**N**H_3_^+^  R466-HN**C**NH_2_^+^  R466-HN**C**NH_2_^+^  R355- HN**C**NH_2_^+^  K356-**N**H_3_^+^  R357- HN**C**NH_2_^+^  E340-**C**NH_2_ | Xyl-1– 2-O-**S**O_3_^-^  Xyl-1– 4-O-**S**O_3_^-^  Xyl-6 – 3-O-**S**O_3_^-^  Xyl-6 – 3-O-**S**O_3_^-^  Xyl-3 – 3-O-**S**O_3_^-^  Xyl-4 – 2-O-**S**O_3_^-^  Xyl-2 – 2-**S**O_3_^-^  Xyl-5 – 3-**S**O_3_^-^  Xyl-4 – 2-**S**O_3_^-^  Xyl-6 – 4-**O**H | 3.7Å (SB)  3.6Å (SB)  5.0 Å (SB)  3.5 Å (HB)  3.8 Å (SB)  4.2Å (SB)  4.8Å (SB)  4.8Å (SB)  4.1Å (HB)  4.0Å (HB) |
|  | **Wuhan Cluster II** | R357-C_α_**N**H R466- HN**C**NH_2_^+^  R355-**C**NH_2_  K356-**N**H_3_^+^  K356-**N**H_3_^+^  K356-**N**H_3_^+^  R346-C_α_**N**H  R346-NH**C**NH_2_^+^ | Xyl-1 – 2-O-**S**O_3_^-^  Xyl-1 – 4-O-**S**O_3_^-^  Xyl-1 – 2-O-**S**O_3_^-^  Xyl-1 – 4-O-**S**O_3_^-^  Xyl-2 – 2-O-**S**O_3_^-^  Xyl-2 – 3-O-**S**O_3_^-^  Xyl-4 – 2-O-**S**O_3_^-^  Xyl-6 – 3-O-**S**O_3_^-^ | 3.6Å (HB)  4.6Å (SB)  3.8Å (HB)  6.4Å (SB)  3.8Å (SB)  5.7Å (SB)  4.2 Å (HB)  4.1 Å (SB/HB) |
|  | **Delta** | R452-NH**C**NH_2_^+^  R346-NH**C**NH_2_^+^  K356-**N**H_3_^+^  K356-**N**H_3_^+^  K356-**N**H_3_^+^  N354-C_α_**N**H  N354-C_α_**N**H  R466-HN**C**NH_2_^+^ | Xyl-1 – 3-O-**S**O_3_^-^  Xyl-4 – 3-O-**S**O_3_^-^  Xyl-1 – 4-O-**S**O_3_^-^  Xyl-2 – 2-O-**S**O_3_^-^  Xyl-2 – 2-O-**S**O_3_^-^  Xyl-4 – 2-O-**S**O_3_^-^  Xyl-4 – 2-O-**S**O_3_^-^  Xyl-4 – 2-O-**S**O_3_^-^ | 5.1Å (SB/HB)  4.1Å (SB)  4.5Å (SB)  5.9Å (SB)  6.0Å (SB)  3.4Å (HB)  4.0Å (HB)  4.1Å (SB/HB) |
|  | **Omicron Cluster I** | K444-**N**H_3_^+^  K444-**N**H_3_^+^  R346-NH**C**NH_2_^+^  R346-NH**C**NH_2_^+^  R346-NH**C**NH_2_^+^  R357-C_α_**N**H  R466- HN**C**NH_2_^+^  R357-C_α_**N**H | Xyl-1 – 2-O-**S**O_3_^-^  Xyl-1 – 3-O-**S**O_3_^-^  Xyl-1 – 3-O-**S**O_3_^-^  Xyl-1 – 3-O-**S**O_3_^-^  Xyl-4 – 2-O-**S**O_3_^-^  Xyl-6 – 2-O-**S**O_3_^-^  Xyl-6 – 2-O-**S**O_3_^-^  Xyl-6 – 3-O-**S**O_3_^-^ | 3.5Å (SB)  4.0Å (SB)  4.7Å (SB)  4.7Å (SB)  5.3Å (SB)  4.2Å (HB)  4.6Å (SB)  3.8Å (HB) |
|  | **Omicron Cluster II** | N450 C_α_**O**NH_2_  R355-C_α_**N**H  R466- HN**C**NH_2_^+^  R466- HN**C**NH_2_^+^  R357- HN**C**NH_2_^+^ | Xyl-6 – 4-**O**H  Xyl-3 – 3-O-**S**O_3_^-^  Xyl-3 – 3-O-**S**O_3_^-^  Xyl-3 – 3-O-**S**O_3_^-^  Xyl-1 – 4-O-**S**O_3_^-^ | 2.7Å (HB)  4.7Å (HB)  5.3Å (SB)  4.5Å (SB/HB)  5.0Å (SB/HB) |

**Supplementary Table 3:** Salt bridges (SB) and hydrogen bonds (HB) established in IAGAIA and PPS docking with the S1-RBD of the Wuhan strain RBD and the Delta and Omicron variants. The pair of atoms on which the distance is calculated are in bold.

**Supplementary Table 4:** Statistical analysis of data presented in Figure 6

| Fig. 6a | Wuhan D614G | Mean±SD: 60.5±5 |  | n=6 |
| --- | --- | --- | --- | --- |
|  | + heparin | Mean±SD: 23.5±2.6 | p<0.0001 | n=6 |
|  | Omicron BA.1 | Mean±SD: 55.2±5 |  | n=6 |
|  | + heparin | Mean±SD: 28.8±4.5 | p<0.0001 | n=6 |
| Fig. 6b | Wuhan D614G | Mean±SD: 60.5±5 |  | n=6 |
|  | + heparin | Mean±SD: 34.5±2.7 | p<0.0001 | n=6 |
|  | Omicron BA.1 | Mean±SD: 55.2±5 |  | n=6 |
|  | + heparin | Mean±SD: 32±2.9 | p=0.0006 | n=6 |
| Fig. 6c | Wuhan D614G | Mean±SD: 60.5±5 |  | n=6 |
|  | + heparin | Mean±SD: 18.3±2.5 | p<0.0001 | n=6 |
|  | Omicron BA.1 | Mean±SD: 55.2±5 |  | n=6 |
|  | + heparin | Mean±SD: 20.8±2 | p<0.0001 | n=6 |
| Fig. 6d | Wuhan D614G | Mean±SD: 60.5±5 |  | n=6 |
|  | + PPS | Mean±SD: 23.7±2.2 | p<0.0001 | n=6 |
|  | Omicron BA.1 | Mean±SD: 55.2±5 |  | n=6 |
|  | + PPS | Mean±SD: 12.7±2.2 | p<0.0001, * p=0.0015 | n=6 |
| Fig. 6e | Wuhan D614G | Mean±SD: 60.5±5 |  | n=6 |
|  | + PPS | Mean±SD: 21.7±3.1 | p<0.0001 | n=6 |
|  | Omicron BA.1 | Mean±SD: 55.2±5 |  | n=6 |
|  | + PPS | Mean±SD: 19.7±3 | p=0.0003 | n=6 |
| Fig. 6f | Wuhan D614G | Mean±SD: 60.5±5 |  | n=6 |
|  | + PPS | Mean±SD: 13.8±3 | p<0.0001 | n=6 |
|  | Omicron BA.1 | Mean±SD: 55.2±5 |  | n=6 |
|  | + PPS | Mean±SD: 5.3±2.6 | p<0.0001, ** p<0.0381 | n=6 |
